# Direct probabilistic quantification of mosaic loss of chromosome Y from sequencing data

**DOI:** 10.64898/2026.06.26.734767

**Authors:** Jhih-Rong Lin, Yoke-Chen Chang, Alexander Maslov, Yinghui Song, Tina Gao, Jidong Shan, David A. Bennett, Sofiya Milman, Nir Barzilai, Jan Vijg, Cristina Montagna, Zhengdong D. Zhang

## Abstract

Loss of chromosome Y (LOY) is the most common aneuploidy in aging men and is increasingly recognized as a marker of aging and genomic instability. Because LOY occurs in mosaic form, its degree reflects the fraction of cells lacking the Y chromosome. Existing SNP-array- and sequencing-based methods rely largely on single genomic features and indirect transformations to estimate this fraction. We developed BaySeq-Y, a Bayesian method that directly estimates LOY mosaicism from sequencing data using VCF files with read depth (DP) and allelic depth (AD). Within a rigorous Bayesian framework, BaySeq-Y integrates complementary LOY-associated genomic features, including decreased read depth and allelic imbalance, and can additionally leverage haplotype phasing to improve precision. In simulations and fluorescence *in situ* hybridization validation (FISH), BaySeq-Y provided accurate estimates and outperformed existing methods. Applications to ROSMAP and GTEx supported its biological relevance through transcriptomic validation, demonstrating its utility for quantifying LOY across diverse sequencing datasets.

## INTRODUCTION

Loss of chromosome Y (LOY) is common in white blood cells of older men and typically occurs in a mosaic form (mLOY), where only a subset of cells within a blood sample have lost the Y chromosome^1^. In peripheral blood, mLOY is generally thought to reflect LOY events in hematopoietic stem and progenitor cells, which then clonally expand and contribute to circulating leukocytes^2,3^. Hematopoietic LOY is rarely detectable before age 50 but becomes increasingly prevalent after age 60^4^. As a distinctive marker of aging and genomic instability, hematopoietic LOY has been associated with increased mortality^4-6^, elevated risk of cancers (including leukemia and non-hematologic cancers)^5,6^, and multiple age-related diseases such as Alzheimer’s Disease^7^ and cardiovascular diseases^8^. Therefore, accurate identification and quantification of LOY have substantial clinical and scientific value.

To date, the most widely used approach for estimating LOY is based on SNP-array genotyping intensities in the non-pseudo autosomal region (non-PAR) of chromosome Y (i.e., mLRR-Y)^1^. Alternatively, LOY can be estimated from the difference between maternal (X-PAR) and paternal (Y-PAR) allelic intensities that are derived from phased SNP-array data as implemented in methods such as PAR-LOY^9^. As mLRR-Y and PAR-LOY use independent signals, a recent GWAS integrated both measures to assess LOY, resulting in a 10% increase in power to detect LOY-associated loci^10^. Similarly, MADloy^11^ utilizes B-allele frequency (BAF) in PAR to capture allelic imbalance to reduce false positive LOY-calls from mLRR-Y. With sequencing data, LOY can also be estimated from the relative sequencing coverage of the Y-non-PAR to autosomal regions^12^ using Control-FREEC^1,13^ or a newer tool such as MosCoverY^14^. In addition to these specialized LOY detection methods, LOY can be estimated using MoChA^15,16^, a general method for detecting mosaic chromosomal alterations. MoChA detects LOY from phased SNP-array or sequencing VCFs based on SNP-array-derived BAF and LRR or sequencing-derived allelic depth in PAR, respectively.

These methods have established the feasibility of using different genomic features to predict LOY across large cohorts. However, there remains an opportunity to further improve LOY estimation by directly quantifying the degree of mosaicism, defined as the fraction of Y-lacking cells. For example, methods such as PAR-LOY provide a dichotomous LOY classification^9^, which is useful for detection but may not fully capture variation in LOY burden across individuals. Other approaches measure the magnitude of LOY-related signal, but converting these signals into the fraction of Y-lacking cells often requires additional transformation and has not always been extensively validated^1,6^. There is also an opportunity to improve estimation by jointly leveraging complementary LOY-associated features, including read-depth and signal-intensity changes in the male-specific region of the chromosome Y and allelic or phased imbalance in the PAR region. Although prior methods have demonstrated the utility of these individual signals, an efficient and flexible framework that integrates them for quantitative LOY estimation could improve both statistical power and biological interpretability. Finally, systematic evaluation against known or orthogonally estimated LOY fractions would further strengthen confidence in quantitative LOY estimates across a broad range of mosaicism levels.

Building on complementary genomic signals associated with LOY, there is an opportunity to develop an approach that directly estimate the fraction of cells lacking the Y chromosome from sequencing data. Motivated by Bayesian approaches for quantifying mosaic aneuploidy in otherwise diploid genomes using sequencing-derived features^17^, we developed BaySeq-Y to directly infer the fraction of Y-lacking cells by jointly modeling multiple LOY-associated sequencing signals. BaySeq-Y is a sequencing-based method that requires only Variant Call Format (VCF) files of whole exome sequencing (WES) or whole genome sequencing (WGS) data containing information of variant read depth (DP) and allelic depth (AD). We systematically evaluated its accuracy using simulation and fluorescence *in situ* hybridization (FISH) and compared its performance with existing methods. We demonstrated its broad applicability and robustness by applying BaySeq-Y to the Religious Orders Study/Memory and Aging Project (ROSMAP)^18^ and Genotype-Tissue Expression (GTEx)^19^ cohorts, for which gene expression data were available for transcriptomic validation. The ability of BaySeq-Y to directly estimate continuous LOY cell fractions provides a unique advantage for studying dose-dependent molecular consequences of LOY. Leveraging this feature, we systematically assessed the relationship between LOY burden and Y chromosome gene expression using GTEx data. Importantly, by directly estimating the fraction of LOY cells within a probabilistic framework, BaySeq-Y enables a more precise and interpretable characterization of mosaicism, facilitating downstream analyses of dose-dependent biological effects.

## RESULTS

### BaySeq-Y integrates complementary genomic signals to directly estimate LOY cell fraction

BaySeq-Y estimates the degree of mLOY for each sample from VCF files derived from WES or WGS data. When read depth and allelic depth information is available for variants in the Y-non-PAR and PAR regions (**Fig. 1A**), respectively, BaySeq-Y can apply a Bayesian model, as defined in Eq. (5) (see **Methods**), that integrates two LOY-associated genomic features: reduced read depth in the Y-non-PAR region and increased deviation of the two BAF peaks of heterozygous variants in PAR from the expected midpoint (see **Methods**). In this model, the fraction of cells lacking chromosome Y is treated as a parameter and estimated from the posterior distribution (**Supplemental Fig. S1**). In addition to point estimates, BaySeq-Y provides full posterior distributions of the LOY fraction, enabling quantification of estimation uncertainty for each sample. When WGS and allelic depth information for phased genotypes in PAR1 are available (**Fig. 1B**), BaySeq-Y can apply a Bayesian model, as defined in Eq. (10), that integrates haplotype-specific genomic features, including decreased allelic depth in the putative paternal PAR1 and haplotype-specific BAF to estimate the LOY fraction. We evaluated the quantitative accuracy of BaySeq-Y using simulations and FISH, for which the true extent of LOY was either known or experimentally determined. In cohort-level applications, we applied BaySeq-Y to the ROSMAP and GTEx cohorts and evaluated the biological validity of the LOY using expression patterns of genes located in the male-specific region of chromosome Y (**Fig. 1C**).

**Figure 1.**
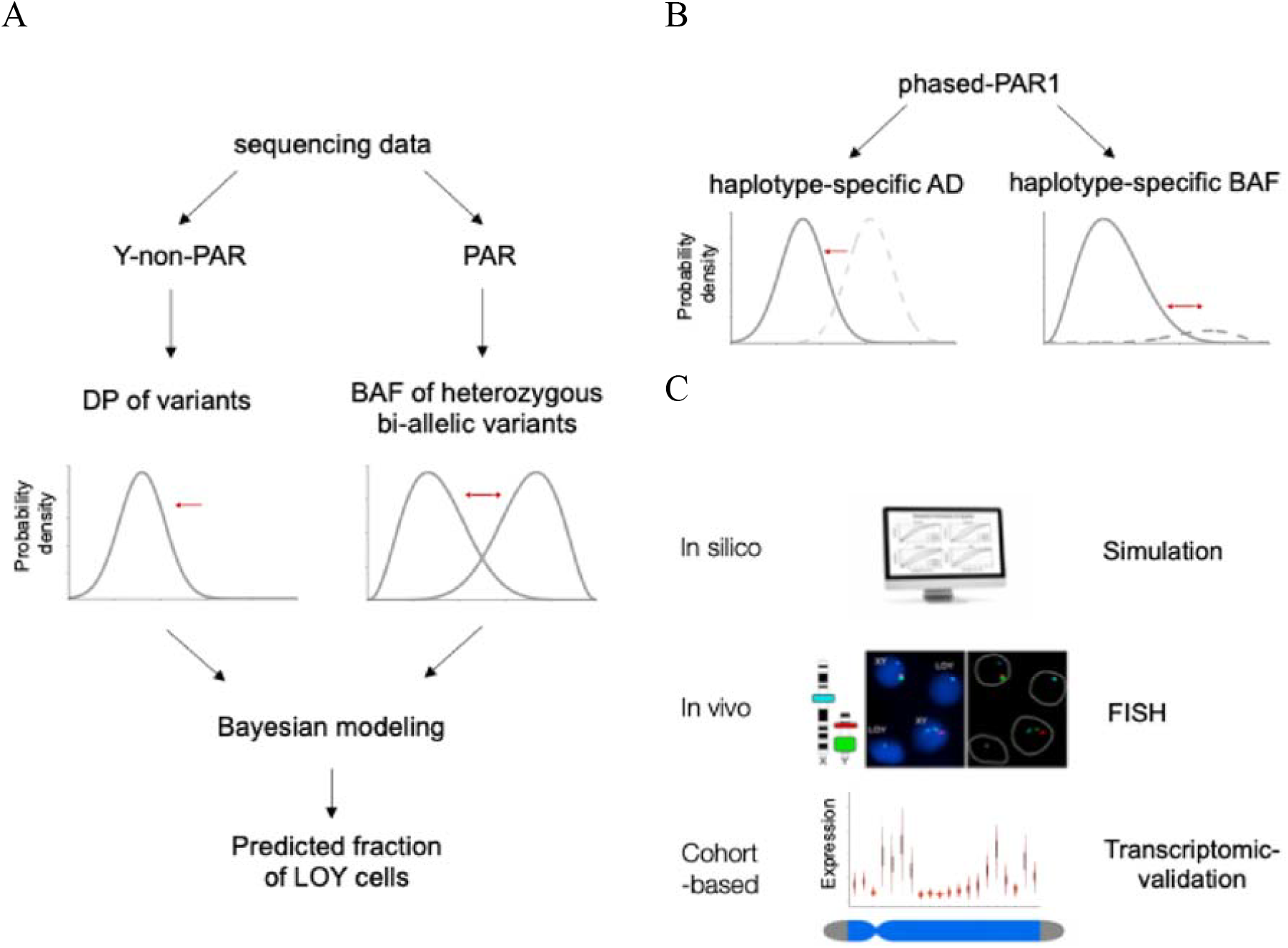
Overview of BaySeq-Y and its performance evaluation. (A) Schematic of LOY estimation by jointly modeling DP in the Y-non-PAR region and BAF in the PAR region. (B) When phased data are available, BaySeq-Y can alternatively estimate LOY using a standalone phasing-based model that incorporates haplotype-specific AD and BAF in the PAR1 region. (C) Strategies used to evaluate BaySeq-Y performance.

Conceptually, BaySeq-Y fundamentally differs from existing approaches in that it directly models the fraction of cells lacking the Y chromosome as a parameter in a unified probabilistic framework, rather than inferring LOY indirectly from individual genomic features. By jointly integrating complementary signals from read depth, allelic imbalance, and phased haplotypes, BaySeq-Y leverages multiple sources of information to improve both accuracy and robustness. This unified modeling strategy enables more precise and interpretable estimation of LOY compared to approaches that rely on single features or post hoc transformations.

### Joint modeling of read depth, allelic imbalance, and phasing enables accurate LOY estimation

To evaluate BaySeq-Y, we first applied it to simulated data under different conditions. These analyses showed that decreased read depth and allelic imbalance provide highly complementary information for LOY estimation: read depth modeling in the Y-non-PAR region is effective at low LOY fractions, whereas BAF modeling in the PAR region is more informative at higher LOY fractions. As expected, sequencing bias reduced the accuracy of θ estimation when only read depth information was used, particularly at higher LOY fractions (**Fig. 2A-B**; **Supplementary Fig. S2**). In contrast, although estimation based solely on the feature of allelic imbalance in the PAR region was unaffected by sequencing bias, it showed limited sensitivity to low-level LOY, likely because the two binomial components become increasingly indistinguishable when their means lie close to the overall mean BAF. By jointly modeling read depth in the Y-non-PAR region and BAF in the PAR region, BaySeq-Y was robust to sequencing bias and outperformed the BAF-modeling approach regardless of whether sequencing bias is high or low. However, its accuracy at low LOY fractions remained suboptimal. Using phased PAR1 data, BaySeq-Y yielded consistently high accuracy across the full range of LOY fractions (**Fig. 2A-B**). We further systematically investigated the performance of BaySeq-Y across randomly assigned LOY fractions between 0 and 0.5, while accounting for different levels of DP bias and phasing quality. BaySeq-Y using phased PAR1 data consistently achieved the best performance (**Fig. 2C** and **Supplementary Fig. S3**), and its accuracy remained stable under different levels of phasing quality. To further investigate the marked improvement in performance and the robustness against phasing errors, we examined the effect of blocking against phasing errors on LOY estimation in BaySeq-Y (phased PAR1). Without partitioning the PAR data into blocks, performance is highly sensitive to the phasing quality (**Fig. 2D**). Dividing the PAR1 region into 20 blocks substantially reduced their impact and maintained stable performance even under a higher phasing error rate by confining the impact of phasing errors to local contaminated blocks.

**Figure 2.**
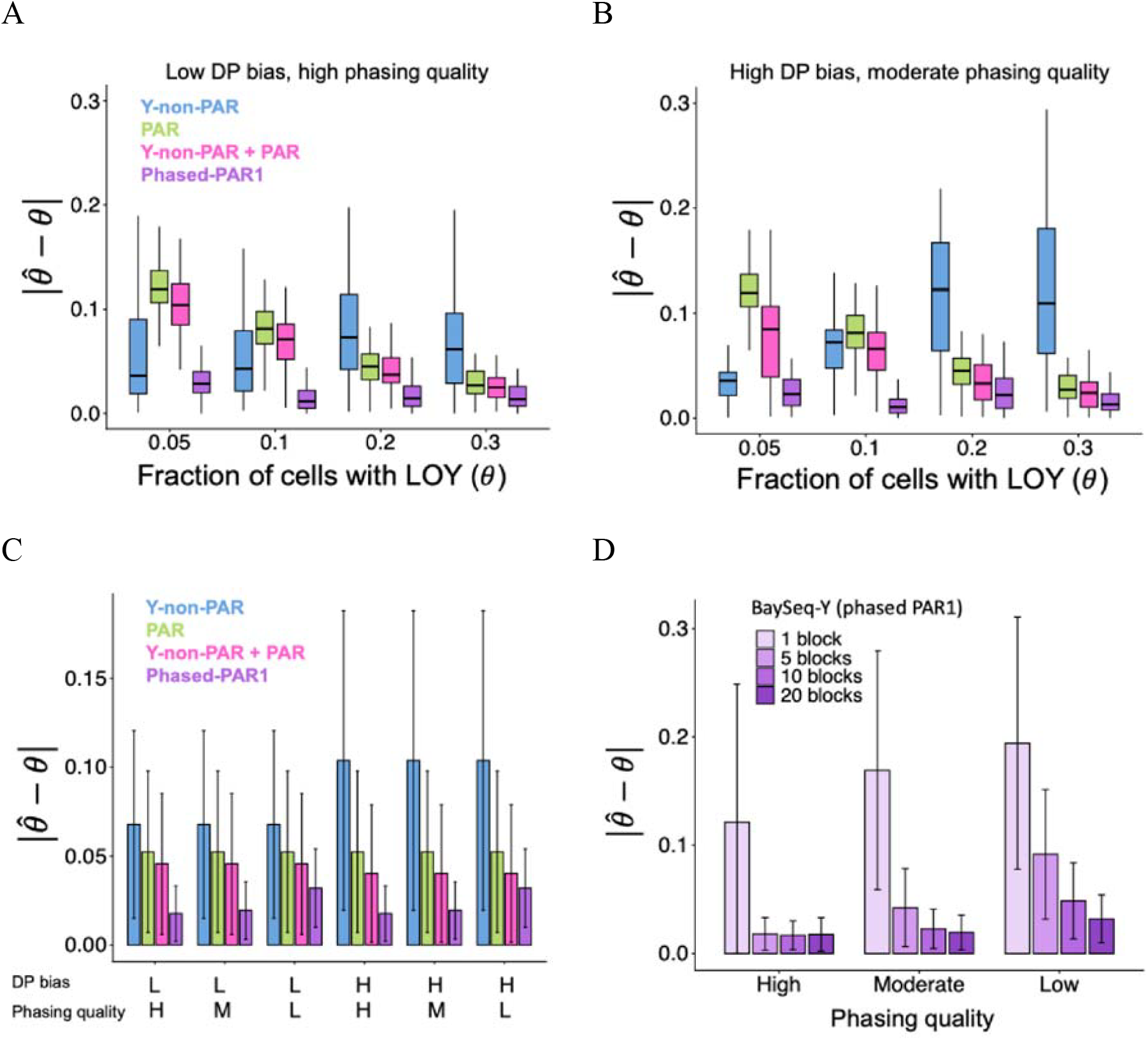
Performance evaluation of BaySeq-Y based on simulation. BaySeq-Y performance using WGS-like data (*n* = 500 variants; see **Methods**) was assessed at four LOY levels with 100 simulated samples per level. Simulations were performed under low DP bias and high phasing quality (A), and high DP bias and moderate phasing quality (B); these simulation settings are defined in **Methods**. Low and high DP bias refer to 10% normalization bias without elevated DP values and 20% normalization bias with 10% randomly elevated DP values, respectively. Elevated DP values were used to mimic the effects of repetitive structures in the Y-non-PAR region. (C) Performance of BaySeq-Y across different input data types. Results are based on 1,000 simulated samples with randomly assigned LOY fractions ranging from 0 to 0.5, evaluated under high (H) and low (L) DP bias and high (H), moderate (M), and low (L) phasing quality settings (see **Methods**). (D) Increasing the number of blocks improves robustness to phasing errors.

### BaySeq-Y remains accurate across sequencing depths and improves detection of low-level LOY

Next, we assessed the effect of sequencing coverage on performance across genome-wide coverages ranging from 10x to 60x. Under the WES setting, where only a limited number of informative variants (*n* = 20 variants) were available, the prediction accuracy of BaySeq-Y (Y-non-PAR + PAR) depended strongly on coverage, with substantial improvement from 10x to 30x (**Supplementary Fig. S4**). In contrast, under the WGS setting, where substantially more informative variants were available (*n* = 500 variants), the prediction accuracy of BaySeq-Y was relatively insensitive to sequencing coverage for BaySeq-Y with and without integration of phasing data (**Fig. 3A-B**). Although performance improved as coverage increased from 10x to 40x, the magnitude of improvement was modest, and both methods still performed reasonably well even at low coverage (10x).

**Figure 3.**
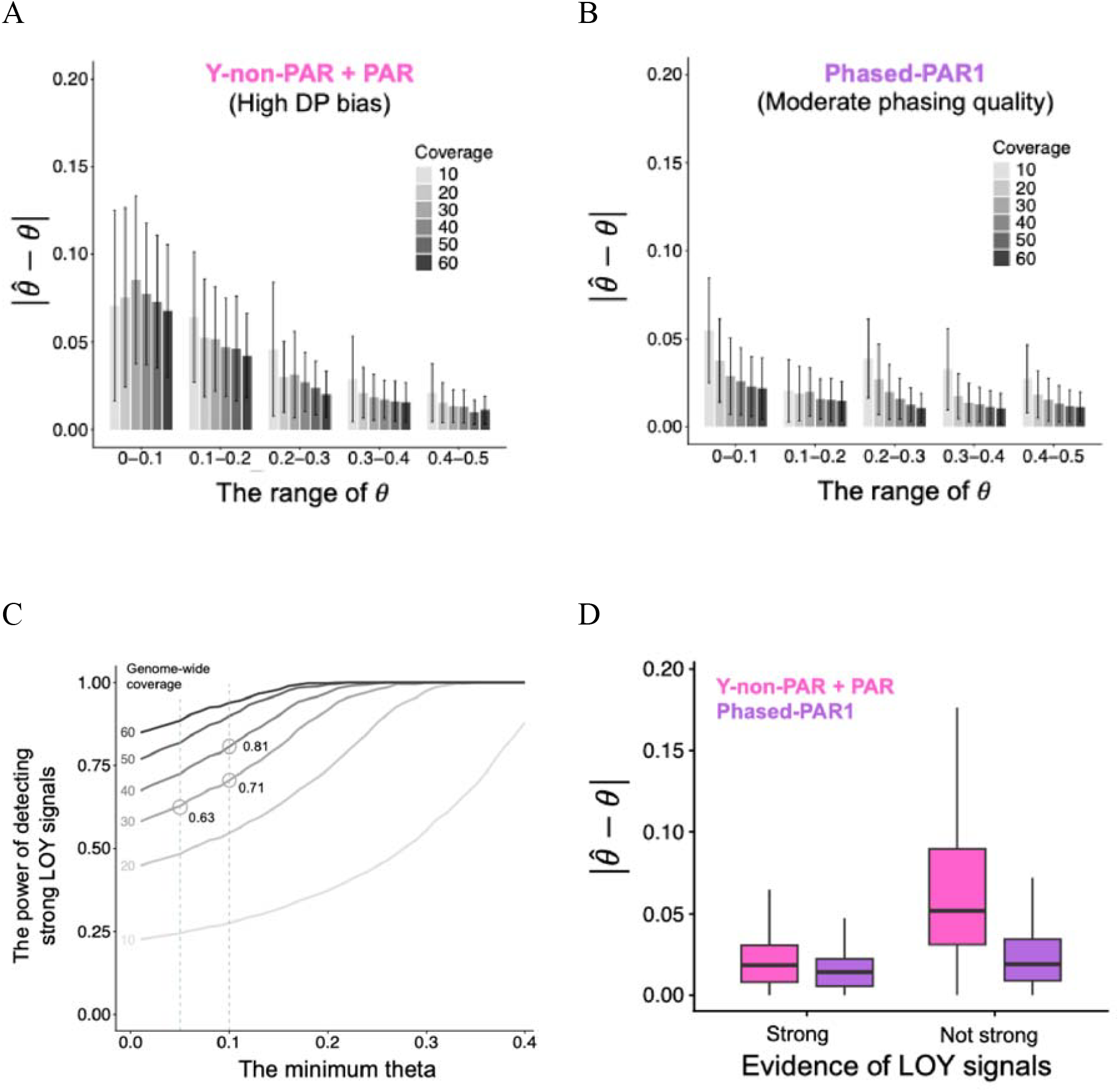
Sequencing coverage versus BaySeq-Y performance. BaySeq-Y performance was evaluated using WGS-like data (*n* = 500 variants) across different sequencing coverages and LOY ranges. All analyses were performed under high DP bias and moderate phasing quality. Performance is shown using (A) combined Y-non-PAR and PAR signals and (B) phased PAR1 signals. Error bars indicate standard deviations. (C) Relationship between WGS coverage and the fraction of subjects with strong LOY signals. Samples were considered to have strong LOY signals when ΔBIC < −10 (see **Methods**). (D) BaySeq-Y performance in samples with and without strong LOY signals.

Sequencing coverage may strengthen LOY signals and improve prediction performance in samples with otherwise weak signals. To investigate this, we used conventional BIC-based model comparison to determine whether a sample showed a strong genomic LOY signal (see **Methods**) and then examined its relationship with sequencing coverage. We observed a strong association between sequencing coverage and the fraction of subjects with strong LOY signals (**Fig. 3C**). Among subjects with at least 10% LOY, based on simulated samples with randomly assigned LOY fractions between 0 and 0.5, the fraction classified as having strong LOY signals increased from 0.71 at 30x coverage to 0.81 at 40x coverage. Next, we investigated the relationship between the detectability of strong LOY signals and the prediction accuracy of BaySeq-Y with and without integration of phasing data. Both methods achieved similarly high accuracy when strong genomic LOY signals were present (**Fig. 3D**). In contrast, BaySeq-Y without phasing integration showed a marked reduction in accuracy among subjects without strong LOY signals, whereas BaySeq-Y with phasing integration maintained high prediction accuracy in these subjects, with only a modest reduction in performance. These results also allow us to assess the minimal detectable level of LOY. Under WGS-like conditions with sufficient variant density, BaySeq-Y with phasing integration was able to reliably detect and estimate LOY at levels as low as ∼10%, with accuracy decreasing gradually below this range as LOY-associated signals become increasingly difficult to distinguish from sequencing noise.

### BaySeq-Y outperforms existing LOY detection methods in simulation and experimental validation

We compared the performance of mLRR-Y, MosCoverY, and MoChA (Three representative methods) and BaySeq-Y in simulations with LOY levels randomly distributed between 0 and 0.5 (**Fig. 4A**). BaySeq-Y with phased PAR1 data outperformed mLRR-Y and MosCoverY in overall accuracy across different levels of DP bias (in Y-non-PAR) and phasing quality (in PAR1). When phasing quality was high, BaySeq-Y achieved performance comparable to MoChA. However, BaySeq-Y maintained stable performance under reasonably degraded phasing quality conditions (i.e., moderate and low phasing quality), whereas MoChA showed a substantial reduction in accuracy. The performance of MosCoverY was highly sensitive to sequencing-related bias. LOY cell fractions converted from mLRR-Y showed the poorest performance, likely reflecting both intrinsic limitations of the underlying signal and reliance on a less well-calibrated empirical conversion formula. Performance across different LOY ranges further showed that BaySeq-Y achieved the highest and most stable accuracy throughout the evaluated LOY range (**Fig. 4B**).

**Figure 4.**
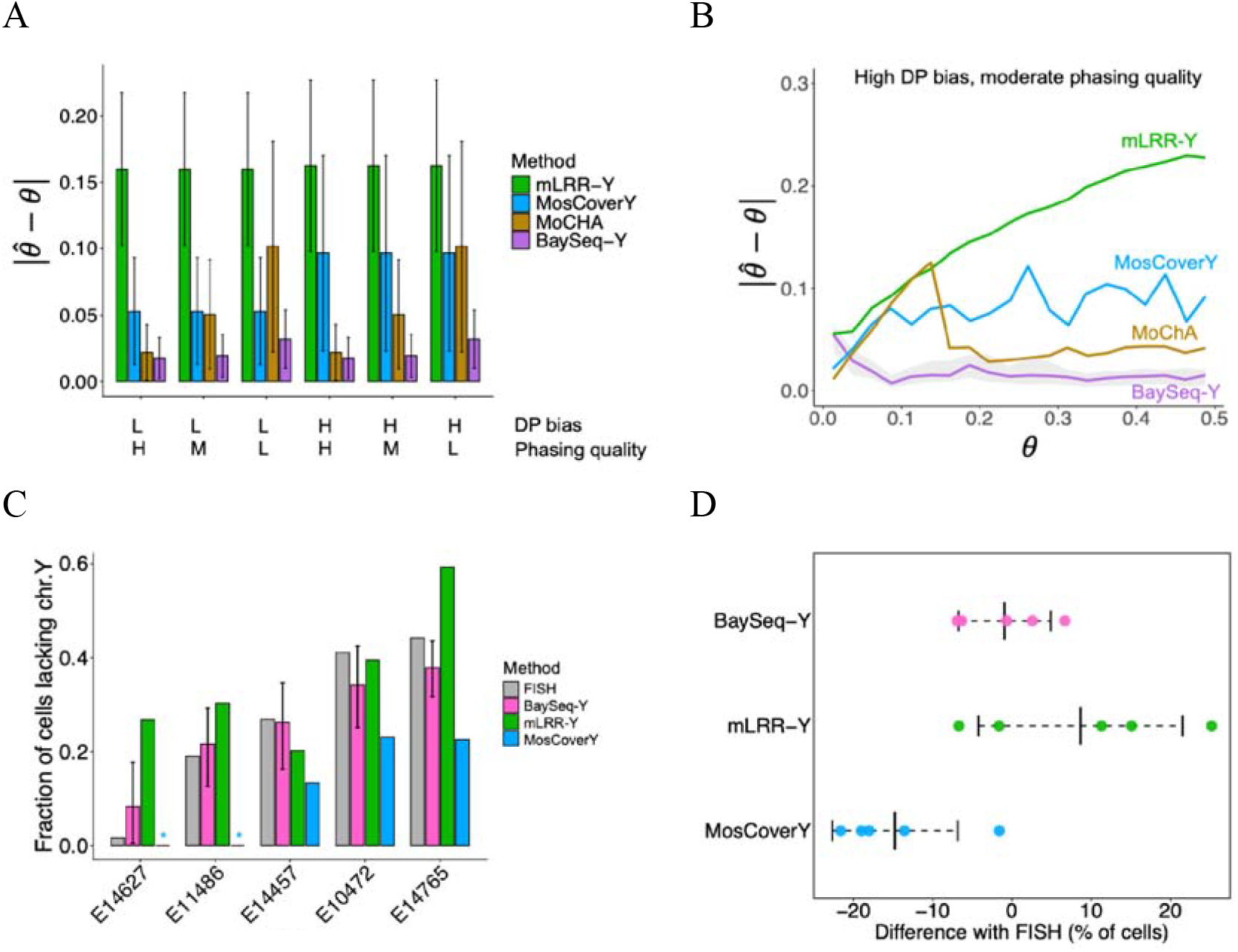
Comparison of BaySeq-Y with other methods. (A) Prediction performance of different methods in simulations of 1,000 samples with LOY fractions randomly assigned between 0 and 0.5. Error bars denote standard deviations. (B) High resolution of prediction performance in simulation at different LOY levels. (C) Experimental validation of different methods using FISH. The asterisk denotes the LOY prediction of 0 for MosCoverY, highlighting reduced sensitivity at low LOY levels. Error bars denote the lower and upper bounds of the 95% confidence interval. (D) The summary of prediction bias from the LOY result based on FISH.

In addition to simulation-based evaluation, experimental validation is essential for assessing the predictions of BaySeq-Y and other methods. We selected five individuals from the Einstein longevity cohort, with ages ranging from 78 to 87 years. The VCF (pVCF) file from the five samples and the corresponding individual BAM files provided by Nonogene were used as input for BaySeq-Y and MosCoverY, respectively. mLRR-Y was calculated from SNP-array signal intensity data. FISH measurements were treated as the reference estimates of the true fraction of cells lacking the Y chromosome, which were 0.016, 0.190, 0.269, 0.411, and 0.442, ordered from low to high (**Fig. 4C**). Although the number of samples is limited, these validation samples span a broad range of LOY levels, enabling evaluation of method performance across both low and high mosaic fractions.

Given WES data, for which statistical phasing of PAR1 is not feasible, we applied BaySeq-Y using Y-non-PAR and PAR data. BaySeq-Y achieved the highest prediction accuracy across the five samples, with an average absolute difference of less than 0.05 (0.046 ± 0.028). The average differences for MosCoverY and converted mLRR-Y were 0.148 and 0.12, respectively (**Fig. 4C-D**). Notably, BaySeq-Y maintained consistent accuracy across the full LOY range. MosCoverY, in contrast, predicted LOY as 0 in the two samples with the lowest true LOY levels, highlighting a fundamental limitation of hard threshold LOY detection methods and raising concerns about low sensitivity or false negatives. These results suggest that BaySeq-Y provides well-calibrated estimates of LOY fraction, with prediction errors that are both small in magnitude and consistent across samples. Although the number of samples used for FISH validation was limited, the selected samples spanned a wide range of LOY levels, providing a stringent test of quantitative accuracy. The consistent agreement between BaySeq-Y predictions and FISH measurements across this range supports the calibration and robustness of the method. Nevertheless, future studies with larger sample sizes will be valuable for further assessing generalizability across cohorts and platforms.

Across both simulation and experimental validation, BaySeq-Y consistently demonstrated superior accuracy and robustness compared to existing methods. In simulations, BaySeq-Y achieved the lowest overall prediction error across the full range of LOY fractions, while maintaining stable performance under varying levels of sequencing bias and phasing quality. In contrast, the higher prediction bias observed for MoChA at an LOY fraction of 0.1 (**Fig. 4B**) was likely because, when the LOY level was low, MoChA did not report any mosaic CNV events, resulting in an estimated LOY fraction of 0. Similarly, our FISH validation suggested that MosCoverY has reduced sensitivity at low LOY levels, often defaulting to zero predictions. In contrast, BaySeq-Y achieved the smallest deviation from experimentally measured LOY fractions across all samples. These differences likely reflect the underlying design of the methods: BaySeq-Y directly models the fraction of LOY cells within a unified probabilistic framework, whereas existing approaches rely on single features and post hoc transformations that are more sensitive to noise and bias. Together, these results indicate that BaySeq-Y provides a more accurate and better calibrated estimate of LOY across diverse conditions.

### Grid-search approximation substantially accelerates BaySeq-Y with minimal loss of accuracy

Under the WES setting, where the number of informative variants is limited, BaySeq-Y with MCMC-based estimation requires approximately 10-20 seconds per sample. In contrast, under the WGS setting, BaySeq-Y requires approximately 5–10 minutes per sample without phasing, while BaySeq-Y using phased data requires approximately 30– 60 minutes per sample, depending on the number of variants. This computational burden may limit its feasibility for large-scale cohort applications. With the fast grid-search implementation, the runtime is reduced to less than 1 minute per sample while maintaining performance comparable to the MCMC-based estimation approach (**Supplementary Fig. S5**).

### BaySeq-Y identifies age-associated LOY and transcriptomic consequences in ROSMAP

We demonstrate the applicability and high accuracy of BaySeq-Y in a real-world application by predicting hematopoietic LOY in ROSMAP. Hematopoietic LOY in ROSMAP has previously been estimated using mLRR-Y from SNP-array data^20^, but SNP-array data are not available for all subjects. Although WGS data from blood are available for 142 male participants, LOY prediction based on sequencing data has not yet been performed because sequencing alignment files are unavailable. Applying BaySeq-Y (phased PAR1 with *m* = 20 blocks) to the VCF file derived from WGS data, we predicted LOY of these 142 males (age: 65.0-95.9; mean ± SD = 78.9 ± 7.4) in ROSMAP. We also applied BaySeq-Y using both PAR and Y non-PAR data and confirmed high consistency between the two approaches (*r* = 0.89) (**Supplementary Fig. S6**), despite their use of different LOY-associated signals. In the following analyses, we focus on results generated by BaySeq-Y using phased PAR1 data. Across the cohort, LOY was observed across a broad range of levels, with approximately 21.1% of individuals showing LOY > 0.1 and 9.2% showing LOY fractions > 0.2, broadly consistent with prior reports of hematopoietic LOY in aging populations^21^, although prevalence estimates vary by age distribution and detection method. LOY showed a modest positive association with age across the cohort. However, this association did not reach statistical significance (*r* = 0.144, *P* = 0.08), possibly reflecting the limited sample size and the exclusion of younger individuals. Among the 142 males, 41 had a diagnosis of Alzheimer’s disease. After adjustment for age, predicted LOY was significantly higher in individuals with Alzheimer’s disease than in those without Alzheimer’s disease (mean LOY: 0.117 vs. 0.078; *P* = 0.036), providing additional support for the biological plausibility of BaySeq-Y predictions^7^.

Among the 142 males, 45 had SNP-array signal data (CEL format), and 20 had bulk RNA-seq data from monocytes (FASTQ format) (**Fig. 5A**). We first compared LOY predicted by BaySeq-Y with converted mLRR-Y derived from the SNP-array signal data (See **Methods**) and observed high concordance (*r* = 0.69) (**Fig. 5B**). The expression of genes in the Y chromosome has been used to detect or infer LOY because of its direct biological relevance^16,22^. Among the 20 individuals with both predicted LOY and monocyte bulk RNA-seq data, one had the highest predicted LOY (= 0.64), whereas the second-highest predicted LOY was 0.28 (**Fig. 5A and 5C**). Notably, examination of the top-expressed MSY genes in these 20 subjects showed that the subject with the highest predicted LOY of 0.64 consistently exhibited a high reduction of expression across MSY genes (**Fig. 5D**). Because VST-transformed expression values are approximately on a log2 scale, the subject with the highest LOY in whole blood (**Fig. 5D**) corresponds to an estimated 80-97% reduction in MSY gene expression relative to baseline in monocytes. The subject with the second-highest predicted LOY (= 0.28) also consistently showed a modest reduction in MSY gene expression, although the effects were more subtle for some MSY genes. Overall, these results are consistent with mLRR-Y, and MSY-gene expression supports the high accuracy of LOY prediction in the ROSMAP application. These results demonstrate that BaySeq-Y captures biologically meaningful variation in LOY that is reflected in downstream gene expression changes.

**Figure 5.**
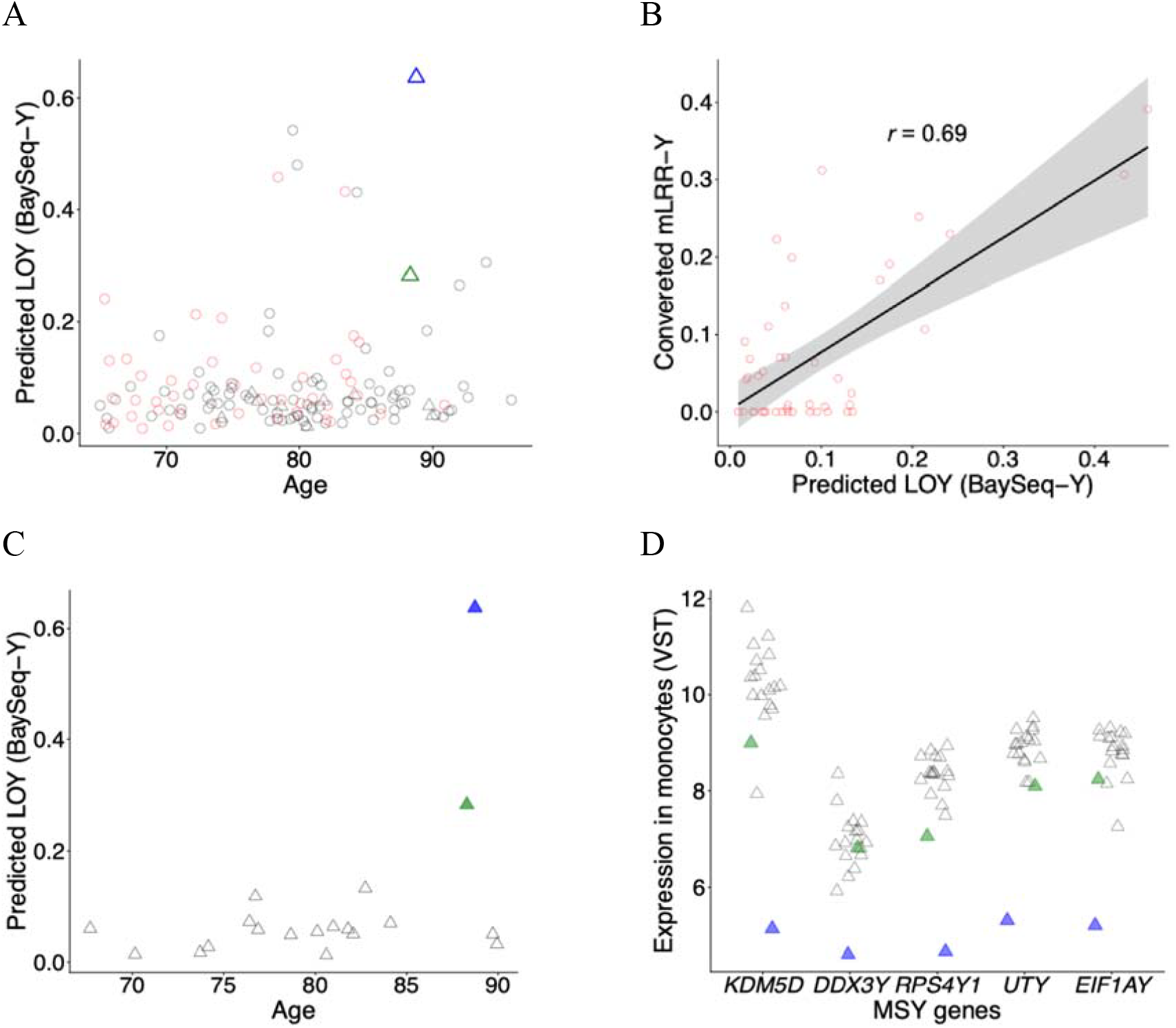
Application of BaySeq-Y in ROSMAP. (A) LOY prediction of 142 subjects using phased PAR1 data from blood WGS. Red denotes 45 subjects with SNP array data available. Triangle denotes 20 subjects with bulk RNA-seq data from monocytes available. (B) Comparison of LOY prediction based on BaySeq-Y and mLRR-Y. (C) LOY prediction of 20 subjects with bulk RNA-seq data from monocytes available. (D) Predicted LOY vs. MSY gene expression. The five most abundant MSY genes in whole blood (GTEx) were shown. Blue and green indicate the samples with the highest and second-highest predicted LOY, respectively, among those with bulk RNA-seq data available.

### BaySeq-Y reveals dose-dependent effects of LOY on Y-chromosome gene expression in GTEx

In this application, we demonstrated the strong applicability of BaySeq-Y in GTEx data, where LOY had not previously been quantified for two main reasons. First, SNP-array data are unavailable, so the conventional SNP-array-based approaches cannot be applied. Second, although WGS data are available, they do not cover the Y chromosome and are provided only as VCF files. To address this, we used BaySeq-Y to integrate phased PAR1 data (*m* = 10 blocks). BaySeq-Y can adapt to data without DP in the male-specific regions in Y chromosomes by leveraging decomposed allelic depth in the PAR region. Unlike the ROSMAP application, we used existing phased data provided by GTEx (v.8) to predict LOY in 377 males. LOY estimates remained low in males younger than 50 years and showed an age-related increase across the cohort after 50 (*r* = 0.20, *P* = 9.3E-05) (**Supplementary Fig. S7**), consistent with the known age-related accumulation of LOY.

We further examined the relationship between LOY and the expression of the five most abundant MSY genes in whole blood among 256 males aged 50 years or older. We observed a clear overall pattern in which individuals with higher LOY levels tended to have lower MSY gene expression (**Fig. 6A**). This observed decrease in gene expression followed a dose-dependent pattern with increasing LOY levels, supporting the quantitative accuracy of BaySeq-Y estimates. Interestingly, the magnitude of expression decline varied across MSY genes and PAR genes (**Fig. 6B**), suggesting that individual genes differ in their sensitivity to LOY. We further systematically evaluated the effects of LOY on gene expression across different LOY levels. 17 out of 21 blood-expressed genes in the PAR and Y-non-PAR regions showed significant susceptibility to LOY (FDR < 0.05) (**Supplementary Table S1**), including five genes reported to be LOY-associated (*TMSB4Y, SLC25A6, CSF2RA, RPS4Y1*, and *CD99*)^16^. Among the remaining 4 genes, *SPRY3* and *VAMP7* did not showed any trend toward decreased expression with increasing LOY (**Fig. 6C-D**). Notably, both genes are located in the PAR2 region and are known to be exclusively transcribed from the X chromosome in males^23^. Some genes appeared more tolerant of low-level LOY (LOY < 0.2), such as *DDX3Y* and *IL3RA* (**Table 1**), whereas *CD99* showed reduced expression even at even lower LOY levels (LOY < 0.1). MSY genes (i.e., *TMSB4Y, UTY, DDX3Y, ZFY*, and *EIF1AY*), whose bulk expression is more strongly driven by monocytes, a cell type known to be enriched for LOY, were more likely to show greater reductions in expression with increasing LOY, although their sensitivities varied. In general, PAR genes were less susceptible to LOY than MSY genes; however, some PAR genes, such as *CSF2RA* and *IL3RA*, still showed substantial expression reduction at high LOY levels. Together, these results demonstrate that BaySeq-Y enables the detection of biologically meaningful, dose-dependent effects of LOY on gene expression at the population level.

**Table 1.** Estimated fold changes in whole-blood gene expression at different LOY levels.

|  | PAR |  |  |  | Y non-PAR |  |  |
| --- | --- | --- | --- | --- | --- | --- | --- |
| LOY | *P2RY8* | *IL3RA* | *CSF2RA* | *CD99* | *UTY* | *DDX3Y* | *KDM5D* |
| 0.1 | 0.93 (0.84-1.03) | 1.05 (0.82-1.34) | 0.95 (0.88-1.04) | 0.92 (0.85-0.99) | 1.00 (0.94-1.06) | 1.07 (0.99-1.15) | 0.97 (0.90-1.04) |
| 0.2 | 0.86 (0.74-1.01) | 1.01 (0.69-1.46) | 0.89 (0.79-1.02) | 0.87 (0.77-0.98) | 0.93 (0.85-1.03) | 1.03 (0.91-1.16) | 0.91 (0.81-1.02) |
| 0.3 | 0.81 (0.68-0.96) | 0.90 (0.60-1.37) | 0.83 (0.72-0.95) | 0.84 (0.74-0.96) | 0.82 (0.74-0.91) | 0.90 (0.79-1.03) | 0.84 (0.74-0.95) |
| 0.4 | 0.76 (0.64-0.91) | 0.77 (0.50-1.17) | 0.75 (0.65-0.87) | 0.83 (0.73-0.95) | 0.69 (0.62-0.77) | 0.74 (0.65-0.85) | 0.76 (0.67-0.86) |
| 0.5 | 0.72 (0.60-0.88) | 0.63 (0.40-0.99) | 0.68 (0.58-0.80) | 0.84 (0.72-0.97) | 0.57 (0.50-0.64) | 0.58 (0.50-0.67) | 0.68 (0.59-0.78) |
| 0.6 | 0.69 (0.54-0.87) | 0.50 (0.28-0.89) | 0.62 (0.51-0.75) | 0.85 (0.71-1.02) | 0.46 (0.39-0.53) | 0.44 (0.37-0.53) | 0.61 (0.51-0.72) |

Note: Values in parentheses indicate 95% confidence intervals.

**Figure 6.**
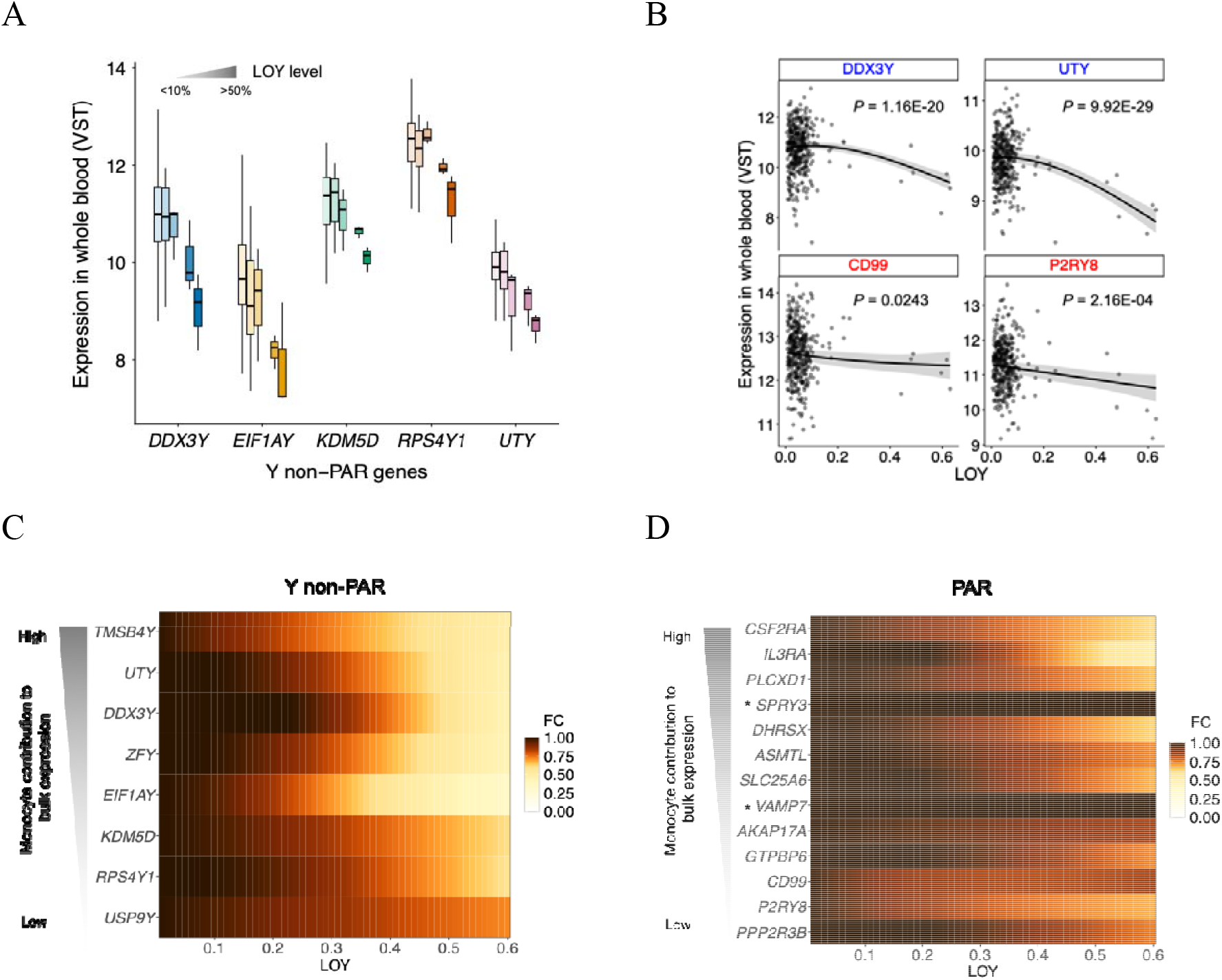
Predicted LOY versus Y-chromosome gene expression using GTEx data. (A) expression of MSY genes among subjects stratified by BaySeq-Y-predicted LOY level. For each gene, the six LOY groups from left to right are: <0.1, 0.1-0.2, 0.2-0.3, 0.3-0.4, 0.4-0.5, and > 0.5. (B) Predicted LOY fraction versus expression of selected genes in the Y-non-PAR region (blue) and PAR region (red). (C, D) Predicted effect of LOY on gene-expression fold change (FC) for Y-non-PAR genes (C) and PAR genes (D). See **Methods** for the monocyte contribution to bulk expression.

## DISCUSSION

Previous LOY methods have been evaluated mainly through correlations with LOY-associated factors such as age, known LOY-associated genetic variants, and smoking status. While these correlations provide indirect support of performance, they do not inform quantitative accuracy or indicate whether the resulting estimates are sufficiently informative for studies of LOY-related effects or phenotypes. Therefore, we employed a multifaceted framework to evaluate BaySeq-Y. Specifically, we used simulation and FISH, for which the true LOY fraction is known or directly measured, to assess quantitative accuracy. Across a wide range of simulated conditions, BaySeq-Y generally achieved high accuracy, with prediction errors typically below 5% (**Fig. 2** and **Supplementary Fig. S2**). The quantitative accuracy observed in simulation was further supported by orthogonal experimental validation using FISH (**4C-D**). We further extended validation through transcriptomic analyses to test whether predicted LOY captures the expected direct biological consequences of chromosome Y loss in cohort data, an important consideration for studies of LOY-related effects or phenotypes. Notably, using LOY estimates from BaySeq-Y, we replicated associations for all five genes in the Y chromosome (PAR and Y-non-PAR) previously reported for MoChA-inferred LOY^16^ and further identified 12 additional genes (**Supplementary Table S1** and **Fig. 6C-D**), despite differences between the datasets.

Our results showed that BaySeq-Y achieved a high accuracy of LOY estimate and outperformed existing methods, primarily for two reasons. First, it is the only method that is specifically designed to directly estimate the fraction of cells lacking the Y chromosome within the model. This estimate is derived within a statistically principled model rather than relying on post-hoc heuristic or empirical transformations. The FISH validation results (**Fig. 4C-D**) suggest that our LOY estimates are well calibrated, whereas other methods that rely on post hoc transformations show substantially different biases across samples with different LOY levels. Second, BaySeq-Y utilizes most of the genomic features informative for LOY, including sequencing depth in both the Y-non-PAR and PAR regions, allelic imbalance in the PAR region, and phasing information. Our simulations showed that integrating these complementary signals substantially improves performance and increases robustness to read-normalization artifacts and sequencing bias (**Fig. 2A-B**).

Phased allelic imbalance in the PAR region can be highly informative for LOY inference and prediction, but proper handling of phasing errors is critical. Previous methods^9,15^ demonstrated this by applying HMM-based frameworks for mosaic copy-number detection that use phased allelic imbalance in the PAR1 region and implicitly account for phasing switch errors across heterozygous sites. However, these frameworks were developed for mosaic copy-number variation rather than for direct estimation of the fraction of LOY cells. In our study, we showed that, within a Bayesian framework specifically designed to estimate the fraction of LOY cells, phasing information can sharpen both read-depth and allelic-imbalance signals to improve LOY prediction (**Fig. 2**). This is because phased read depth avoids the normalization bias affecting Y-non-PAR depth, while even small amounts of LOY produce a clear shift in the mean of the dominant binomial component away from the paternal haplotype. The shift remains detectable because the binomial component arising from potential maternal haplotypes due to phasing errors is relatively small. We adopted three procedures that effectively mitigate the impact of phasing errors. First, in both the DP and BAF models, we used a mixture model to account for the component arising from maternal haplotypes due to phasing errors. Second, we divided the PAR region into n blocks and modeled them jointly with a shared LOY estimate but block-specific parameters, which helps localize the impact of phase-switch errors. Third, we introduced a hidden variable to identify contaminated blocks and down-weight their influence on LOY estimation. Simulation results showed that these procedures effectively mitigate the impact of phasing switch errors, allowing BaySeq-Y to perform well even at reasonably high phasing switch error rates (**Fig. 2C-D**).

Our Bayesian framework was inspired by MADSEQ, a previously developed method for detecting mCAs^17^. Although MADSEQ established a Bayesian strategy for estimating the cellular fraction of mosaic chromosomal alterations from allelic imbalance and depth signals, it was limited to autosomes and chromosome X in females under an assumption of chromosomal diploidy and required sequence alignment data in addition to variant calls. MADSEQ has not been widely used for mCA calling, likely because it does not leverage phased allelic imbalance, which has been shown to be highly informative, while DP signals in diploid genomes are often noisy and have limited sensitivity for detecting aneuploidy events. Here, we showed that a Bayesian framework can effectively incorporate LOY-related genomic features for accurate LOY prediction. This is mainly for two reasons. First, unlike depth signals in diploid genomes, depth in the haploid Y-non-PAR region provides a much sharper signal for aneuploidy event (i.e., LOY). Second, we incorporated phasing information, which sharpens the read-depth and allelic-imbalance signals within the phased component, thereby substantially improving prediction, particularly at low to moderate LOY levels. More broadly, BaySeq-Y represents a shift from heuristic or feature-specific LOY estimation toward a unified probabilistic framework that directly models the underlying biological quantity of interest. By explicitly parameterizing the fraction of cells with LOY and integrating multiple complementary genomic signals, BaySeq-Y avoids reliance on empirical transformations and provides estimates that are both quantitatively interpretable and statistically grounded. This conceptual framework may be broadly applicable to the study of other forms of somatic mosaicism where multiple noisy signals must be integrated to infer cellular heterogeneity.

Although high levels of LOY are relatively easy to detect, the ability of a method to accurately estimate low-level LOY, and the importance of doing so, have been underappreciated. This problem has been largely overlooked because validation of LOY prediction at low levels has rarely been conducted, and the significance of low-level LOY has remained unclear. The fundamental challenge in predicting low-level LOY is that LOY-related signals are often weak and difficult to distinguish from noise arising from multiple sources, including normalization bias, alignment bias in sequencing reads, and phasing errors. We showed that, by integrating multiple LOY-related features within a statistical framework, low-level LOY, which usually lacks strong signals, can still be estimated with reasonable accuracy (**Fig. 2A-C**). By contrast, threshold-based methods that assume zero LOY when strong signals are lacking, such as MosCoverY, can produce substantial bias at low LOY levels. In the FISH validation, the two samples with the lowest LOY levels were both predicted as 0 by MosCoverY; one was close to the observed value (0.016), whereas the other substantially underestimated the observed LOY level (0.19). Using GTEx data, we showed that bulk expression changes in some MSY or PAR genes (e.g., *CD99*) can be detected in whole blood at LOY levels as low as 0.1 (**Table 1**). These findings suggest that BaySeq-Y is capable of detecting low-level LOY with reasonable accuracy and underscore the importance of accurate LOY estimation, even at low levels, for studying the biological effects and associated phenotypes of LOY.

Using a method specifically designed to estimate the fraction of cells with LOY, we performed, to our knowledge, the first analysis of the effects of LOY on gene expression in GTEx data. Although our inference relied only on genomic features in the PAR region, it is reasonable to interpret the observed aneuploidy signal as LOY, because other sex-chromosome aneuploidies in males, such as loss of X or gain of X or Y, are extremely rare. Our results provide at least three insights. First, expression of Y-chromosomal genes in blood appears to differ in its sensitivity to LOY (**Fig. 6**). This likely reflects differences in cell-type contribution to bulk expression, enrichment of LOY in specific cell types, and gene-specific regulatory mechanisms. Second, we found that several PAR genes with important immune-related functions, including *IL3RA, CSF2RA*, and *P2RY8*, can show substantially reduced expression at higher levels of LOY, for example around 0.3-0.5 (**Fig. 6D** and **Table 1**). Third, although some PAR genes, such as *SPRY3* and *VAMP7*, may not be actively transcribed from the Y chromosome in males, our results suggest that at least 9 PAR genes show active transcription from the Y chromosome in blood cells (**Supplementary Table S1** and **Fig. 6D**).

Compared with other sequencing-based methods that depend on sequence alignment files^13,14^, which are not always available, BaySeq-Y requires only VCF files with read depth (DP) and allele depth (AD) information. These files are routinely generated in most sequencing studies and are substantially easier to manage in large cohorts. More generally, BaySeq-Y illustrates how quantitative information retained in standard VCF files can be leveraged to infer somatic mosaicism without requiring access to large sequencing alignment files. Unlike methods relying on accurate phasing information^14^, BaySeq-Y integrates it as an optional feature to improve the performance. Moreover, BaySeq-Y is applicable to both WES and WGS without requiring external control samples, unlike some LOY callers that are largely limited to WGS^13,15^ or require matched controls for WES^13^ data. This broader applicability and scalability can expand opportunities for LOY research, particularly in settings where alignment files are unavailable, only WES data are available, or reliable phasing data cannot be obtained. An additional advantage of BaySeq-Y is that it provides posterior distributions for LOY estimates, allowing uncertainty to be explicitly quantified. This is particularly valuable for samples with low LOY levels, where distinguishing true signal from noise is inherently challenging.

## METHODS

### Study participants

We applied BaySeq-Y to two cohorts with available WGS data: ROS and MAP, and GTEx. ROS and MAP are aging cohort studies designed to study aging, Alzheimer’s disease and related traits. They have a large common core of identical data elements collected by the same staff, allowing efficient merging of data into the ROSMAP dataset^24^. This cohort is particularly useful for LOY analysis because it is enriched for older individuals and includes multiple genomic and transcriptomic data modalities^18^. In addition to WGS data, ROSMAP includes SNP-array data, which enabled comparison with mLRR-Y, as well as bulk gene-expression data from monocytes, which we used for transcriptomic validation. In ROSMAP, 1,196 samples were whole-genome sequenced, among which 391 were generated from blood-derived DNA. Among these blood-derived samples, 142 were male and used in our analyses of LOY prediction. The sequencing coverage of this dataset was approximately 30x.

GTEx is a project designed to study gene expression across multiple human tissues, including whole blood, with matched WGS data available for many donors. This cohort provided an opportunity not only to perform transcriptomic validation of BaySeq-Y predictions, but also to systematically investigate the relationship between LOY and expression of Y chromosome genes in whole blood. We used GTEx v8, in which both WGS and whole-blood RNA-seq data were available for 377 male donors, including 256 donors aged 50 years or older. The WGS data were generated at approximately 30x coverage.

### Bayesian modeling of LOY using sequencing data

When phased data in the PAR region are unavailable, BaySeq-Y can detect and quantify LOY using two Bayesian models: one for sequencing depth in the Y-non-PAR region and the other for BAF at heterozygous sites in the PAR region. It integrates independent LOY signals from these two regions by maximizing the combined likelihood of the LOY fraction – the fraction of cells with LOY – across both models (**Fig. 1A**). Unlike other sequencing-based approaches that require sequencing alignment files^1,13,14,17^, BaySeq-Y requires only WES/WGS VCF files containing AD and, optionally, read depth (DP), which is equal to the sum of allelic depths. Although VCF files are primarily used to represent germline variant calls, they also retain quantitative read-level information, such as AD and DP, at each variant site. These measurements reflect the underlying mixture of cell populations in the sample and therefore capture signals of somatic mosaicism, including mosaic loss of chromosome Y.

#### Bayesian modeling of Y-non-PAR

For each sample to be predicted, we first processed Y-non-PAR variants and X-non-PAR variants for the non-PAR read-depth component of the model. Assuming short-read sequencing data, we retained only variants separated by more than 300 bp to reduce local dependence among sampled read depths, consistent with the independence assumption in our Bayesian model. We then accounted for GC-related coverage bias^25^ by matching X non-PAR variants to the GC-content distribution of Y non-PAR variants. Specifically, for each analyzed Y-non-PAR or X-non-PAR variant, local GC content was calculated as the fraction of G and C bases within a 100-bp window centered on the variant position in the reference genome. Variants in the Y-non-PAR and X-non-PAR regions were separately assigned to 10 GC-content bins, according to the intervals of width 0.1 (0-0.1, 0.1-0.2, …, 0.9-1.0). For each sample, we then randomly selected X-non-PAR variants from the corresponding GC-content bins so that the sampled X-non-PAR variants matched the GC-bin distribution of the retained Y-non-PAR variants. The mean read depth of these GC-matched X-non-PAR variants was used to approximate the expected Y-non-PAR read depth in the absence of LOY.

In our Bayesian modeling for LOY, *θ* denotes the fraction of cells with LOY. The expected mean read depth of variants in the Y-non-PAR region, *m*_DP_, can be represented as:

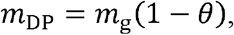

where *m*_g_ represents the expected mean variant read depth in the absence of LOY and was approximated using the mean read depth of the GC-matched X-non-PAR variants. We modeled the observed read depth using a negative binomial distribution to account for overdispersion^17^:

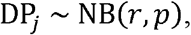

where *p* is parameterized to match the expected mean *m*_DP_,

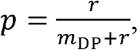

and *r* controls dispersion, which was assigned a weakly informative Gamma prior^17^,

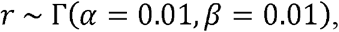

Accordingly, for *n* variants, the likelihood function for observed read depths is:

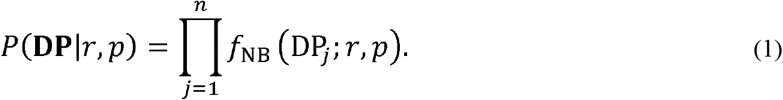

#### Bayesian modeling of PAR

Given the sequencing data for an individual, we constructed a second Bayesian model for LOY using allelic read depth at heterozygous bi-allelic variants in the PAR region. We define BAF as the AD of alternative alleles divided by the total read depth of a variant. For a diploid chromosome, BAF is expected to be centered near a midpoint around 0.5, but it shifts toward 1 and 0 in the presence of mosaic monosomy^17^. BAF in the PAR region is expected to show a similar deviation when LOY is present. Accordingly, we modeled the observed AD of heterozygous variants in the PAR region using a hierarchical two-component beta-binomial (BB) mixture model:

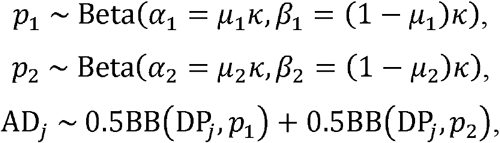

where DP_*j*_ is the read depth of variant *j*. The parameters *μ*_1_ and *μ*_2_ represent the expected BAF means given the midpoint *m* and the fraction of cells with LOY, *θ*:

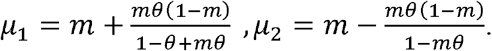

The midpoint *m* is estimated as the median BAF for all heterozygous bi-allelic variants in autosomal regions for the subject. The concentration parameter *κ*, which controls the variance of BAF, is assigned as a Gamma prior:

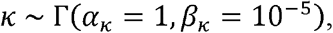

where the shape and rate parameters correspond to a weakly informative prior allowing large variance. Accordingly, for *n* variants, the likelihood function for observing allelic depths is:

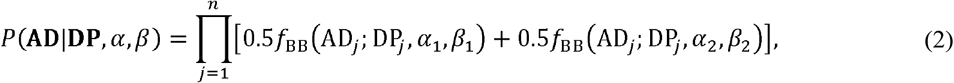

where *f*_BB_ denotes the beta-binomial probability mass function.

#### Block-wise joint modeling for robust LOY estimation

Sequencing-derived DP and BAF signals may be affected by unmodeled components, such as technical noise, repeat-enriched regions, mappability differences, or other sources of systematic deviation, raising concerns about model misspecification. When the number of informative variants is small, as in WES data, sporadic deviations in DP or BAF can often be accommodated by flexible dispersion parameters and may have limited influence on the estimate of the LOY fraction *θ*. In contrast, when many variants are available, as in WGS data, repeated or directionally consistent deviations in DP or BAF are less likely to be fully absorbed by a single global variance structure and may instead distort the estimated *θ*. To reduce this sensitivity, we partitioned the observed variant information into m blocks and applied the same DP model for Y-non-PAR variants and AD model for PAR variants within each block. The LOY fraction *θ* was shared across all blocks, whereas nuisance parameters were estimated independently for each block. This block-wise formulation relaxes the assumption that all variants share a single nuisance structure, allowing heterogeneous deviations in depth or allelic imbalance to be partially absorbed within blocks rather than being forced into the shared LOY estimate. Accordingly, the implementation of this block-wise joint Bayesian modeling for Y-non-PAR read depth and PAR allelic depth can be formulated, respectively, as:

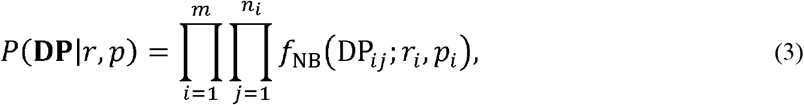

and

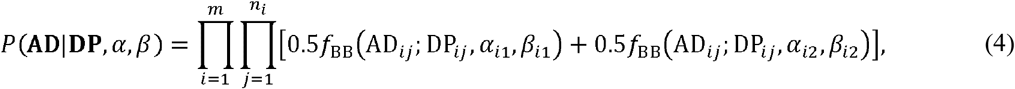

where *i* indexes blocks, *j* indexes variants within block *i*, and *n*_*i*_ is the number of variants in block *i*.

#### Model combination and parameter estimation

Given the likelihood function for the observed variant read depth in the Y-non-PAR region, *P*(**DP**|*p, r*), and the likelihood function for the observed allelic depth in the PAR region, *P*(**AD**|**DP**, *α, β*), we defined the combined likelihood function as the product of the two likelihood components:

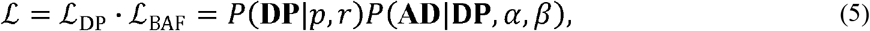

corresponding to the product of Eq. (1) and Eq. (2) for WES data and the product of Eq. (3) and Eq. (4) for WGS data. Given this likelihood function and the prior distribution for the model parameters, the fraction of cells with LOY, *θ*, was estimated from its posterior distribution using Markov Chain Monte Carlo (MCMC) sampling. MCMC sampling was performed in R using JAGS through the rjags package, with 10,000 burn-in iterations followed by 10,000 sampling iterations across three Markov chains.

#### Determination of strong LOY signals for a sample

We constructed a Bayesian statistical model to infer the fraction of cells with LOY. This framework also allowed us to assess how well the LOY model fits the data relative to a null model without LOY. A better fit of the LOY model indicates stronger evidence for LOY in the sample. To quantify this evidence, we used the Bayesian Information Criterion (BIC) to compare the LOY model with the null model. The null model used the same Bayesian modeling framework for both Y-non-PAR and PAR regions, but assumed that there was no LOY. For the Y non-PAR region, the likelihood function was the same as in Eq. (1), except that the parameter *p* was defined as:

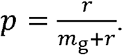

Under the assumption of no LOY, the observed allele depth in the PAR region can be simplified as a single beta-binomial distribution:

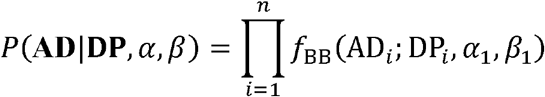

Where the parameters are defined as above, except that *μ*_*1*_ is set as m since *θ* = 0. To compare the LOY model with the null model, we monitored the deviance during posterior sampling for each model and used the minimum sampled deviance, *D*_min_, as an approximation to 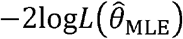 The BIC was then calculated as:

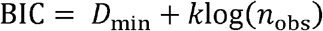

where *k* is the number of estimated model parameters, and *n*_obs_ is the number of observations included in the likelihood. Evidence for LOY was quantified by comparing the BIC of the LOY model with that of the null model. We defined a sample as having strong evidence of LOY if ΔBIC < –10, where ΔBIC = BIC_LOY_ – BIC_null._

### Integrating phasing information in PAR for better LOY estimation

#### Statistical basis for modeling phased data

While long-read sequencing provides direct physical phasing information, phased genotypes can also be statistically inferred from short-read WGS data using tools such as SHAPEIT^26^. When phased allele-depth information is available, we can construct an enhanced Bayesian model using phased heterozygous variants in PAR1. This model extends the read-depth component described in Eq. (1) and the BAF component described in Eq. (2) by incorporating haplotype-specific allele-depth information.

For each sample, allele depths are available for the two phased haplotypes in PAR1. We denote these as AD_S_ and AD_L_, corresponding to the haplotypes with smaller and larger mean allele depth, respectively. However, it is not appropriate to directly assume that *AD*_*s*_ always represents the paternal PAR haplotype, because under no or low LOY, sequencing noise, and phasing errors, the paternal haplotype may not consistently have lower observed allele depth. Therefore, we introduced a latent indicator variable, *v*, to represent whether AD_S_ corresponds to the paternal haplotype:

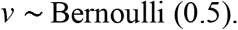

For variant *j*, the allele depth assigned to the paternal PAR haplotype can be written as:

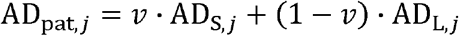

Similarly, the allele depth assigned to the maternal PAR haplotype was defined as:

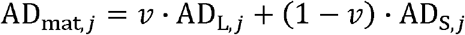

This formula allows the paternal/maternal haplotype assignment to be inferred during posterior sampling rather than being fixed a priori based on the observed mean allele depth. Because statistical phasing is imperfect, the allele depth assigned to the putative paternal PAR haplotype may include reads originating from the maternal haplotype. To account for this, we modeled the observed allele depth in the putative paternal PAR haplotype, AD_pat_, using a two-component mixture model:

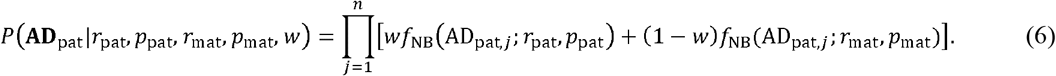

Here, *w* represents the proportion of correctly phased allele-depth observations and was assigned the prior distribution:

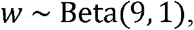

which favors correct phasing while allowing tolerance for phasing switch errors. The expected mean allele depth of the paternal PAR haplotype was defined as:

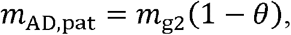

where *m*_g2_ represents the expected mean allele depth of paternal PAR haplotype in the absence of LOY was approximated as:

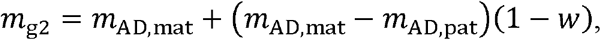

where *m*_AD,mat_ and *m*_AD,pat_ denote the observed mean allele depths in the putative maternal and paternal PAR, respectively. The negative-binomial probability parameters for the paternal and maternal components were then defined as:

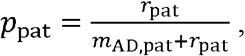

and

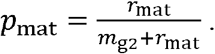

The dispersion parameters for the paternal and maternal components were constrained to be equal, *r*_pat_ = *r*_mat_, and the shared dispersion parameter was assigned a weakly informative prior, consistent with the prior used in the Y-non-PAR model:

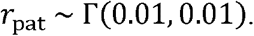

Unlike the Y-non-PAR read-depth model, which requires GC-matched variants in X-non-PAR for read depth normalization. This procedure was not required for the PAR haplotype-specific allele-depth model because the putative paternal and maternal allele depths are measured at the same variants and are therefore matched for sequence context.

When phasing information was incorporated into the beta-binomial framework, the observed allele depth aligned to the putative paternal haplotype was modeled as a mixture of two beta-binomial components:

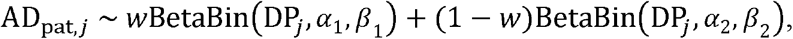

where DP_*j*_ is the total read at variant *j, w* is the proportion of corrected phased observations, and (1-*w*) represents the contribution from phasing-error-affected observations. The parameters *α*_1_, *β*_1_, *α*_2_, and *β*_2_ were defined as described above. The corresponding likelihood is:

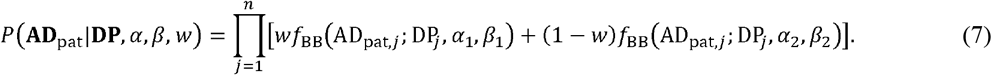

#### Modeling strategies to mitigate phasing errors

Statistical phasing is not expected to be perfectly accurate, and even a single switch error can obscure or reverse the read-depth difference between the two phased haplotypes. In addition to the mixture component described above, which accounts for phasing switch errors at the variant level, we applied two additional modeling strategies to reduce the influence of phasing-error-affected regions. First, PAR1 variants were ordered by genomic coordinate and divided into *m* blocks, each containing an approximately equal number of variants. We then constructed block-specific sub-models that shared a common LOY fraction estimate, *θ*, while allowing all other model parameters to vary independently across blocks. This structure allows the model to borrow information across PAR1 through the shared LOY parameter while accommodating local heterogeneity caused by phasing errors or regional noise. Second, for each block *i*, we introduced a latent contamination variable, *z*_*i*_, to represent whether the block was affected by phasing error. The prior distribution of *z*_*i*_ was specified as:

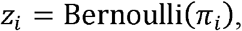

with

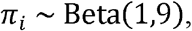

reflecting the prior expectation that most individual blocks are locally consistent in haplotype phase, particularly when the number of blocks is sufficiently large (*m* ≥ 10). The model components described above were used for clean blocks (*z*_*i*_ = 0). For contaminated blocks (*z*_*i*_ = 1), we defined nuisance components designed to capture flattened or distorted allele-depth signals caused by phasing switch errors while reducing their dependence on the shared LOY fraction *θ*. Specifically, in the nuisance components, *p*_*pat*_ and *p*_*mat*_ in Eq. (3),*p*_1_ and *p*_2_ in Eq.(4), and *w* in both equations were assigned weakly informative uniform priors:

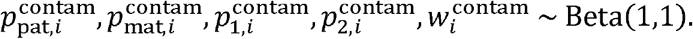

For each block, we then defined effective parameters according to its inferred contamination status:

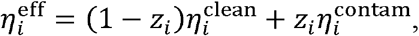

where *η*_*i*_ denotes any block-specific parameter entering the likelihood. These effective parameters were substituted into the likelihood functions in Eq. (6) and (7). For the negative binomial model in Eq. (6), the block-specific likelihood becomes:

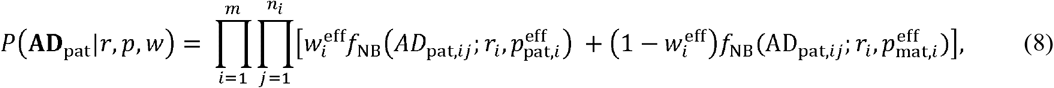

where *i* indexes blocks, *j* indexes variants within block *i*, and *n*_*i*_ is the number of variants in block *i*. For the beta-binomial model in Eq. (7), the block-specific likelihood becomes:

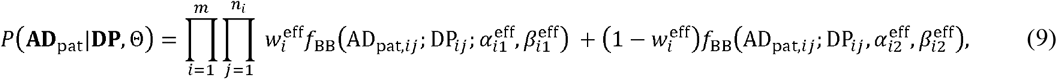

where

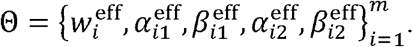

The rationale for this formulation is that blocks inferred to have a higher probability of contamination contribute less directly to the estimation of the shared LOY fraction *θ*. Instead, their distorted signal is absorbed by nuisance parameters, thereby reducing the influence of phasing-error–affected blocks on the LOY estimate.

The combined likelihood function for the full model of BaySeq-Y with integration of phasing data (BaySeq-Y(P)) was defined as the product of the two likelihood components described in Eqs. (8) and (9):

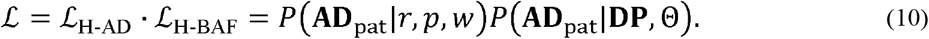

*L*_H-AD_ denotes the haplotype-specific allele-depth likelihood, and *L*_H-BAF_ denotes the likelihood of haplotype-specific BAF, which is defined as AD of paternal/maternal alleles divided by the total read depth of a variant.

### Grid-search estimation of LOY fraction

The original MCMC implementation provides a rigorous Bayesian approach for estimating the LOY cell fraction, but it can be computationally intensive. As a faster alternative, we implemented a grid-search/profile-likelihood approximation for phased PAR allele-depth data. Instead of continuously sampling over the LOY fraction, the fast implementation evaluates a predefined grid of candidate LOY fractions, by default from 0 to 1 in increments of 0.001. For each candidate LOY fraction, the expected haplotype-specific allele imbalance is calculated using the baseline BAF, and the likelihood of the observed haplotype-specific allele depths is evaluated across genomic blocks. Nuisance parameters are approximated by finite grids or mixture likelihoods rather than sampled by MCMC. They include the beta-binomial allele-count concentration parameter (*κ*), the block phase-orientation parameter assigning phased haplotypes to the maternal/X-PAR high-BAF versus paternal/Y-PAR low-BAF bands, the within-block correct-phase fraction for clean blocks (*w*), and block-level contamination modeled through a global contamination rate (*π*).

For each block and LOY grid value, the clean-block likelihood is marginalized over combinations of κ, phase orientation, and clean-block *w*. A separate contaminated-block likelihood is precomputed for each block, allowing contaminated blocks to follow flexible allele-balance patterns that are not constrained by the global LOY fraction. The clean and contamination likelihoods are then combined across a grid of possible global contamination rates, with a conservative Beta (1,9) prior on *π* and a cap on per-block log-Bayes-factor contributions to reduce the influence of extreme blocks. Finally, to regularize estimation in the presence of weak or noisy LOY signals, a beta prior, Beta (1,9), on the LOY fraction is applied to obtain a posterior profile, from which the reported LOY fraction is the grid value with the highest posterior profile probability.

In this study, we will use the fast grid-search estimation method to evaluate BaySeq-Y with phased PAR1 data in simulation. For the two real applications of BaySeq-Y in the ROSMAP and GTEx cohorts, we will use MCMC-derived estimates.

### Performance evaluation

We applied three complementary strategies to evaluate the prediction performance of BaySeq-Y: Simulation, FISH, and transcriptomic validation.

#### Simulation

To evaluate the performance of BaySeq-Y through simulation, we first defined the genome-wide sequencing coverage as *c*. Most simulations were performed under *c* = 30x, reflecting a commonly used WGS or WES coverage setting. To evaluate the effect of sequencing coverage on BaySeq-Y performance, we additionally performed simulations under other coverage values. For each subject, we simulated read depth for *n* variants in the Y-non-PAR region, BAF data for *n* variants in the PAR region, and phased BAF data for *n* variants in the PAR region. We set *n* = 20 and *n* = 500 to represent WES-like and WGS-like scenarios, respectively. Phased BAF data only apply to the WGS-like scenario.

Depending on the simulated LOY fraction *θ*, read-depth data in the Y-non-PAR region were generated from a negative binomial distribution with mean

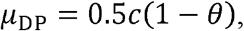

and variance

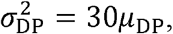

based on empirical observations. The negative binomial parameters were then defined as

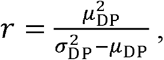

and

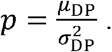

Thus, read depth was simulated as

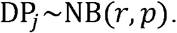

We introduced two types of sequencing bias into the Y non-PAR read-depth simulation: normalization bias and repeat-related alignment bias^27^. Normalization bias was modeled at two levels, low and high, whereas repeat-related alignment bias was modeled as either present or absent. To simulate normalization bias, the expected mean read depth of the X non-PAR region, *m*_X_, was generated from a normal distribution and then rounded to the nearest integer:

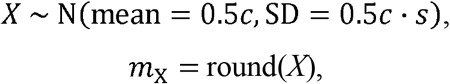

where *s* controls the magnitude of normalization bias. We set *s* = 0.1 for low normalization bias and *s* = 0.2 for high normalization bias. When repeat-related alignment bias was present, 10% of variant read depths were randomly selected and replaced with values generated from an independent negative-binomial distribution with mean = 2*μ*_DP_ and variance = 30×2*μ*_DP_. This procedure was designed to mimic locally elevated read depth caused by repeat-related alignment artifacts in the Y-non-PAR region^27^. Across our simulations, we defined low and high DP bias as 10% normalization bias without randomly elevated DP values and 20% normalization bias with 10% randomly elevated DP values, respectively.

To generate the BAF data in the PAR region, we first simulated read-depth data for PAR variants and then generated allele-depth data for each variant. Read depth in the PAR region was generated from a negative binomial distribution with mean

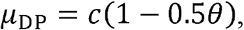

and variance

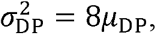

based on empirical observations from the autosomal variants. For each variant, read depth was generated from the negative binomial distribution as described above. For each variant, we then simulated the allele depth of the alternative allele conditional on the simulated read depth. Let DP_*j*_ denote the simulated read depth for variant *j*, and let *m* = 0.5 denote the expected BAF in the absence of LOY. Under LOY fraction *θ*, the two expected BAF values were defined as:

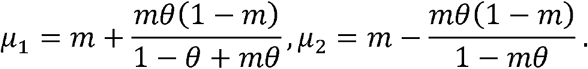

The alternative-allele depth was then generated from an equal-weight mixture of two binomial distributions:

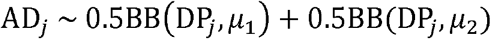

In our simulations, we introduced BAF noise by randomly selecting 5% of PAR variants and replacing their simulated alternative-allele depth with a value generated from a random BAF. Specifically, for selected variants, the alternative-allele depth was set to

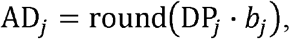

where

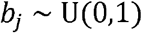

This procedure was used to mimic variants with highly noisy or uninformative allelic imbalance signals.

To generate phased BAF data, we first generated read depth data of PAR variants as described above. For each variant *j*, let DP_*j*_ denote the simulated total read depth. We then simulated haplotype-specific allele depths by assigning reads to the two phased haplotypes according to the expected BAF under LOY. Specifically, with *m* = 0.5, the expected allele fraction for the maternal haplotype was defined as

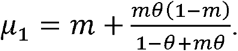

The read depth assigned to this haplotype was then generated as:

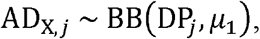

and the read depth assigned to the paternal haplotype was defined as the remaining read depth:

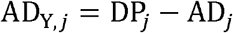

Next, we introduced BAF noise and phasing errors. To simulate BAF noise, we randomly selected 5% of variants. For each selected variant, the haplotype-specific allele depths were replaced with noisy values while preserving the direction of imbalance between the two haplotypes. Specifically, if AD_X,*j*_ ≥ AD_Y,*j*_, the noisy allele depths were generated as

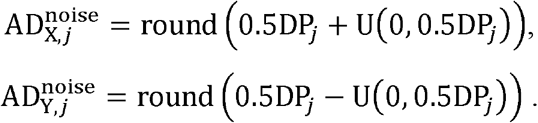

If AD_X,*j*_ *<* AD_Y,*j*_, the same procedure was applied in the opposite direction. This procedure introduced noisy variation in haplotype-specific allele depths while maintaining the original orientation of the larger and smaller haplotype signals. We simulated data under three tiers of phasing quality: high, moderate, and low. For high-quality phasing, we introduced a switch error rate of 0.2%, consistent with trio-based validation of large-scale UK Biobank data^21^, together with a 1% local error rate to represent isolated phasing errors. For moderate- and low-quality phasing, we used switch error rates of 1% and 3%, respectively, and local error rates of 5% and 10%, respectively.

To simulate the performance of mLRR-Y, we assumed that probe-level R ratios in the Y-non-PAR region followed a gamma distribution with mean = 1 – *θ*, where *θ* is the simulated LOY fraction. Based on empirical observations, the variance was set to (0.1 + 0.5*θ*)(1 – *θ*). The corresponding shape and rate parameters were derived from the specified mean and variance. For each simulated sample, we generated 1,000 probe-level R ratios, calculated the mean log R ratio, mLRR-Y, and converted it to the estimated fraction of cells (CF) with LOY using an empirical formula^1^:

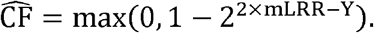

To simulate the performance of MosCoverY, we first generated reference data by simulating read depth values in the Y non-PAR and autosomes for 10,000 samples without LOY, using the negative-binomial framework described above. From this reference population, the population median of normalized chromosome Y coverage, denoted *c*Y_ref_, was calculated once. Evaluated samples were then simulated under the specified LOY fractions using the same read-depth framework. The two types of sequencing bias were introduced using a similar approach described above. According to the formula in MosCoverY, for each evaluated sample, the scaled normalized chromosome Y coverage was calculated as

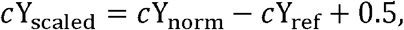

and *c*Y_ref_ was converted to the estimated fraction of cells with LOY as

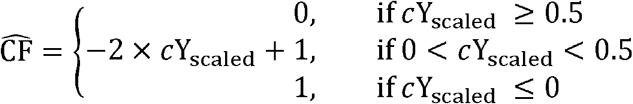

We ran MoChA using the bcftools plugin to call LOY from simulated VCF data in the PAR region. The simulated VCFs were designed to match the simulation setup used for BaySeq-Y, consisting of 500 variants evenly spaced across PAR1. Allelic depths were generated under the same distributional assumptions across simulated LOY fractions, with the same phasing-quality settings: high, moderate, and low. A MoChA LOY call was defined as a mosaic CNV event in PAR1 with length > 2 Mb. The LOY fraction was estimated from the MoChA BAF deviation estimate, denoted as *b*_dev_, using the formula 4*b*_dev_ /(1 + 2*b*_dev_).

#### *Invivo* validation

Five random PBMC samples from the EINSTEIN longevity cohort^28^, for which SNP-array signal data were available, were selected for in vivo validation of BaySeq-Y and comparison with representative SNP-array and sequencing-based methods. Each of the five PBMC samples was divided into two aliquots, which were processed separately for DNA extraction for WES and for FISH analysis.

Genomic DNA was extracted from peripheral blood mononuclear cells (PBMCs) using the Zymo Quick-DNA Miniprep Plus Kit (Zymo Research) according to the manufacturer’s instructions. Briefly, approximately 1-1.5×10^6^ PBMCs per sample were lysed, and genomic DNA was purified using silica spin-column technology. DNA was eluted in the supplied elution buffer and quantified using a Qubit fluorometer (Thermo Fisher Scientific). DNA purity was assessed by NanoDrop spectrophotometry (Thermo Fisher Scientific), and DNA integrity was evaluated using the TapeStation system (Agilent Technologies). Samples meeting quality-control requirements were used for whole-exome sequencing.

Whole-exome sequencing (WES) was performed by Novogene (Sacramento, CA, USA). Genomic DNA was randomly fragmented to approximately 180–280 bp, followed by end repair, A-tailing, and ligation of Illumina sequencing adapters. Adapter-ligated fragments were PCR amplified, size selected, and purified. Exonic regions were enriched by hybridization capture using biotin-labeled probes and streptavidin-coated magnetic beads, followed by additional PCR amplification of captured libraries. Library quality was assessed using Qubit fluorometric quantification, real-time PCR, and Bioanalyzer analysis of fragment size distribution. Qualified libraries were pooled and sequenced on an Illumina platform using 150-bp paired-end reads, with approximately 6 Gb of sequencing data generated per sample.

Raw sequencing reads underwent quality control to remove adapter-contaminated reads, reads containing excessive ambiguous bases (>10%), and reads with a high proportion of low-quality bases (>50%). Clean reads were aligned to the human reference genome using Burrows-Wheeler Aligner (BWA), followed by sorting with SAMtools and duplicate marking with Picard. Single-nucleotide variants (SNVs) and small insertions/deletions (indels) were identified using the Genome Analysis Toolkit (GATK), and variants were annotated using ANNOVAR. Standard sequencing and alignment quality metrics, including mapping rate, duplication rate, target-region coverage, sequencing depth, base quality scores (Q20/Q30), and sequencing error rate, were evaluated by Novogene as part of their routine quality-control workflow. Because the resulting VCF files retain quantitative read-level information, including allelic depth (AD) and total read depth (DP), they served as the primary input for BaySeq-Y estimation of mosaic LOY.

Fluorescence *in situ* hybridization (FISH) was performed on PBMC aliquots from the same five individuals used for whole-exome sequencing. PBMCs were fixed in methanol:acetic acid fixative (3:1) and prepared on glass slides for interphase FISH analysis. Three-color FISH was performed using Vysis CEP Y (DYZ3) SpectrumOrange Probe, Vysis CEP Y (DYZ1) SpectrumGreen Probe, and Vysis CEP X (DXZ1) SpectrumAqua Probe. Hybridization was performed according to the manufacturer’s protocol with minor laboratory-specific modifications. Briefly, slides were aged at 95°C for 5 min, denatured at 75°C for 3 min, and hybridized with the probe mixture at 37°C overnight. Slides were then washed under post-hybridization conditions using 0.4× SSC/0.3% NP-40 at 74°C for 3 min, followed by 2× SSC/0.1% NP-40 at room temperature for 3 min. Slides were then dehydrated through a 70%, 90%, and 100% ethanol series for 3 min each at room temperature, air-dried, counterstained with DAPI, and mounted using Vectashield (Vector Laboratories) mounting medium.

Image acquisition was performed using the BioView imaging system with unbiased automated image acquisition, targeting approximately 500 nuclei per sample. For each field, 15 focal planes were acquired and merged for signal evaluation. FISH scoring was performed blinded to BaySeq-Y predictions and other computational LOY estimates. Only intact, non-overlapping interphase nuclei with interpretable hybridization signals were included in the analysis. Nuclei with ambiguous, overlapping, truncated, or weak hybridization signals were excluded from scoring.

FISH signal patterns were recorded for the DYZ1 SpectrumGreen, DYZ3 SpectrumOrange, and DXZ1 SpectrumAqua probes. A nucleus was classified as Y-retaining when a CEP X signal and Y-chromosome probe signals were detected. A nucleus was classified as LOY when a CEP X signal was present but both Y-chromosome probe signals, DYZ1 and DYZ3, were absent. The FISH-based LOY fraction was calculated as the number of LOY nuclei divided by the total number of scorable nuclei for each sample. Across the five samples, 448, 511, 516, 545, and 487 nuclei were scored, respectively, for a total of 2,507 nuclei. The corresponding FISH-based LOY fractions were 1.6%, 19.0%, 44.2%, 41.1%, and 26.9%, respectively.

#### Transcriptomic validation

For ROSMAP data, the bulk gene expression RNA-seq data for monocytes (SynID: syn22024496) are provided in FASTQ format. RNA-seq reads were aligned to the hg38 reference genome using STAR^29^, with the genome index generated from the reference genome FASTA and GENCODE v48 primary assembly annotation. Coordinate-sorted BAM files were produced for each paired-end sample, and gene-level read counts were subsequently obtained using featureCounts^30^. The count matrix was then processed in R by retaining gene identifiers and sample count columns, standardizing sample names, and converting the matrix into GCT format. For GTEx, gene-level read counts from bulk whole-blood RNA-seq were obtained from the GTEx website in GCT format (gene_reads_2017-06-05_v8_whole_blood.gct). The same downstream processing workflow was applied to both the ROSMAP and GTEx datasets. Briefly, the GCT-formatted gene count matrix was imported into DESeq2 in R, followed by size-factor normalization to account for differences in sequencing depth across samples and variance-stabilizing transformation (VST) to generate normalized expression values for downstream expression-based analyses.

### Data preprocessing

#### Phasing of WGS data in ROSMAP

Common variants in the PAR1 region of ROSMAP WGS data were phased with SHAPEIT5 (phase_common_static)^26^ with a European 1000 Genomes (Freeze 3) chromosome X as the reference panel and the hg19/b37 PAR1 genetic map file from the official website.

#### LOY estimation in mLRR-Y in ROSMAP

The goal was to compare BaySeq-Y prediction with mLRR-Y estimates in samples for which both blood-derived WGS and SNP-array data were available. SNP-array signal intensity data were not available in the ROSMAP dataset cataloged in Synapse. Upon request, the Rush team provided CEL files for 1,280 subjects. By matching genotypes from these CEL files to the SNP-array data available in ROSMAP, we identified 45 male individuals who also had blood-derived WGS. Probe signal intensity (LRR) in the male-specific region of chromosome Y was calculated for the 45 male samples as log_2_(*R*_observed_/*R*_expected_), where *R*_expected_ was estimated as a weighted linear combination of probe signal intensities from 50 male offspring samples in the HapMap 3 panel^31^. The weights were obtained by linearly regressing the probe intensities of the HapMap reference samples against those of the query sample.

### Modeling the relationship between LOY and Y-linked/PAR gene expression

Twenty-one Y-non-PAR and PAR genes expressed in whole blood in GTEx, defined as median TPM > 10, were included in the analysis of the GTEx cohort. For each gene, the association between predicted LOY and VST-transformed expression was modeled using a natural spline term for predicted LOY with 2 degrees of freedom, adjusting for age, top five genetic PCs, the top 20 inferred covariates, PCR status, and sequencing platform. To calculate the predicted fold change, expression was estimated across LOY values, and fold change was calculated relative to the predicted expression at LOY = 0. Because the VST-transformed expression is approximately on the log2 scale, fold change was calculated by exponentiating the predicted expression contrast using base 2.

The monocyte contribution to bulk expression was approximated for each selected gene by dividing the summed expression in classical monocytes by the summed expression across all PBMCs among male subjects aged 50–70 years, using the scRNA-seq PBMC dataset (SynID: 49637038).

## CODE AVAILABILITY

All code required to reproduce the results reported in this study and to apply BaySeq-Y to new datasets is available at GitHub (https://github.com/zdz-lab/BaySeq-Y). BaySeq-Y is implemented in R using the rjags package for Markov chain Monte Carlo (MCMC)-based Bayesian inference. A grid-search implementation for rapid LOY estimation from phased PAR1 data is also provided.

## DATA AVAILABILITY STATEMENT

ROSMAP data are available through the AD Knowledge Portal (https://adknowledgeportal.org). GTEx whole-blood bulk RNA-seq data (gene_reads_2017-06-05_v8_whole_blood.gct) are publicly available through the GTEx Portal□(https://gtexportal.org). Individual-level GTEx whole-genome sequencing data analyzed in this study are available through controlled access from dbGaP (accession phs000424).

## ACKNOWLEDGMENTS

The results reported in this study are based in part on data obtained from the AD Knowledge Portal□. ROSMAP data were generated and distributed by the Rush Alzheimer’s Disease Center, Rush University Medical Center, Chicago. We thank Shinya Tasaki for providing access to the SNP-array CEL files. The Genotype-Tissue Expression (GTEx) Project was supported by the Common Fund of the Office of the Director of the National Institutes of Health and by NCI, NHGRI, NHLBI, NIDA, NIMH, and NINDS. GTEx data used in this study were obtained from the GTEx Portal□ (November 22, 2025) and from dbGaP (accession phs000424; September 16, 2020). We thank the study participants and staff of the Rush Alzheimer’s Disease Center. This research was funded by U19 AG056278, R56 AG088624, R21 AG093047, and R01 AG044829 from the National Institute on Aging, National Institutes of Health. The content is solely the responsibility of the authors and does not necessarily represent the official views of the National Institutes of Health.

## Notes

### Competing Interest Statement

The authors have declared no competing interest.

