## Supplemental Data for "Direct probabilistic quantification of mosaic loss of chromosome Y from sequencing data"

### Supplementary Figures

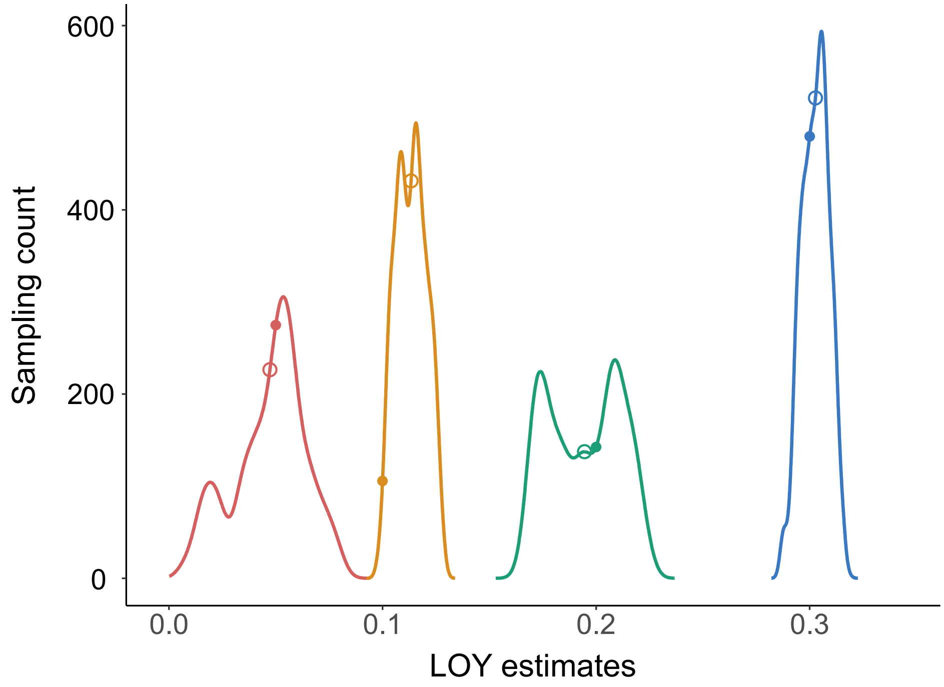


**Supplementary Figure S1.** Posterior distributions of LOY estimates. The curves show the posterior distributions of 10,000 sampled values of $\theta$ (LOY estimate) after burn-in during MCMC sampling. These distributions were generated from simulated data for four subjects with true LOY values of 0.05, 0.1, 0.2, and 0.3, respectively (solid dots). Hollow dots indicate the median posterior LOY estimate, which was reported as the predicted LOY.

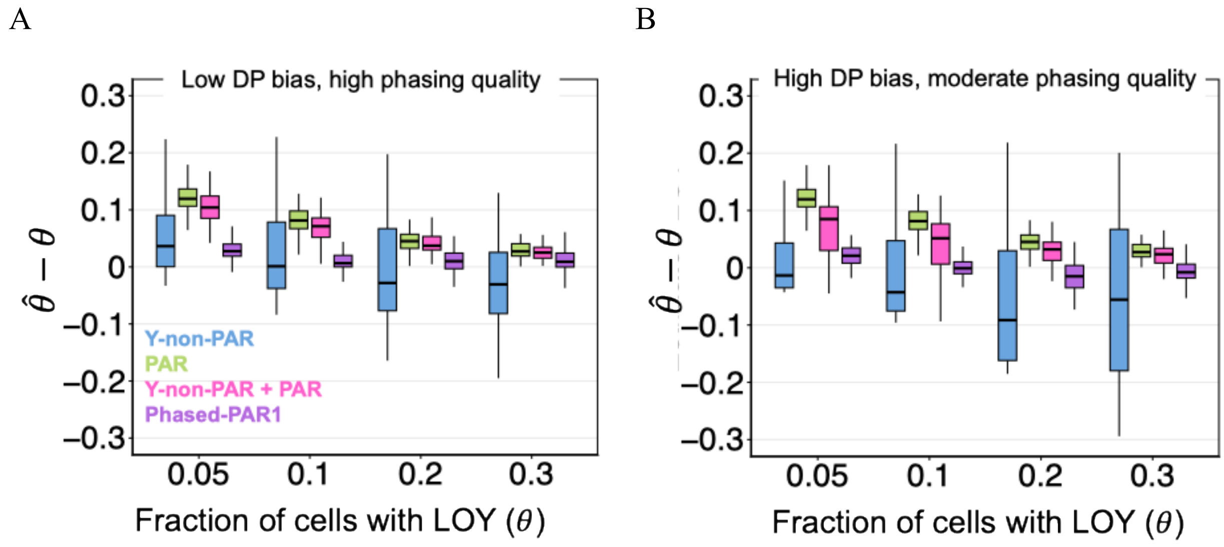


**Supplementary Figure S2.** Evaluation of prediction error of BaySeq-Y based on simulation. BaySeq-Y performance using WGS-like data ($n$ = 500 variants, see **Methods**) was evaluated at four LOY levels (100 simulated samples each) using simulations under low DP bias and high phasing quality (A), and high DP bias and moderate phasing quality (B).

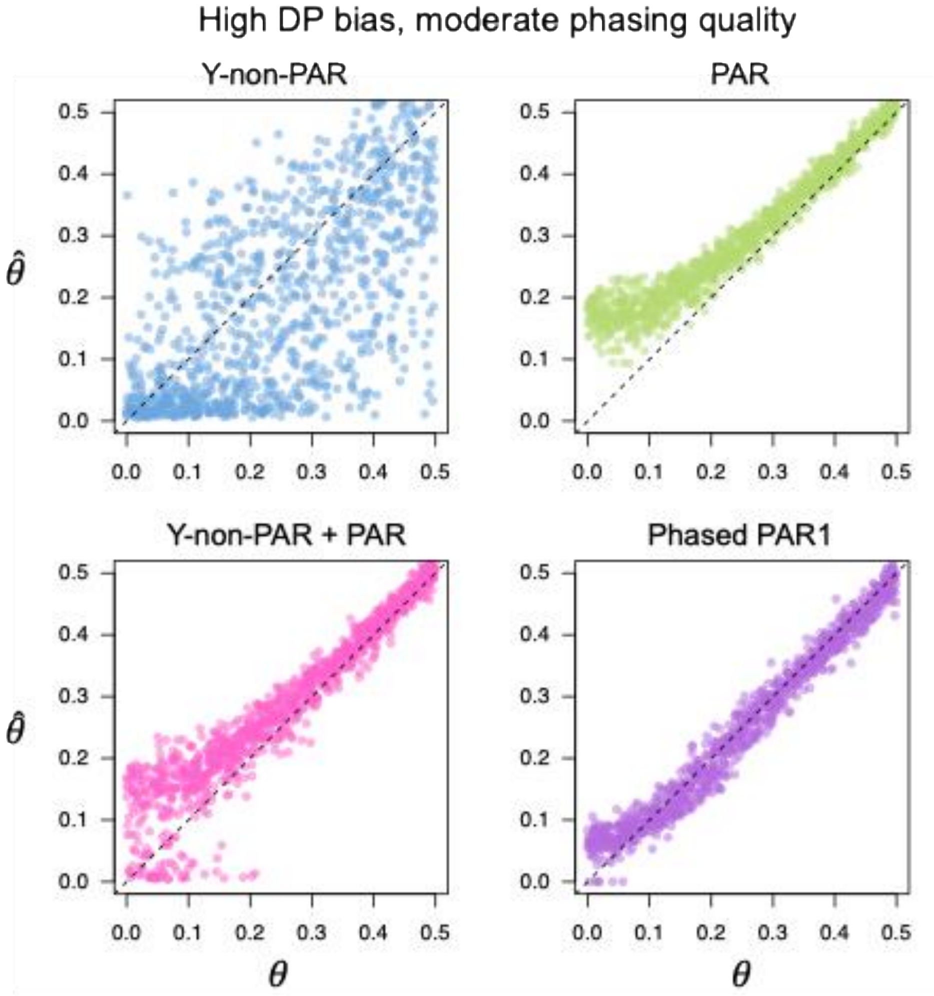


**Supplementary Figure S3.** Predicted LOY fractions were compared with true LOY fractions across 1,000 simulated samples with randomly assigned LOY fractions between 0 and 0.5 under high DP bias and moderate phasing quality.

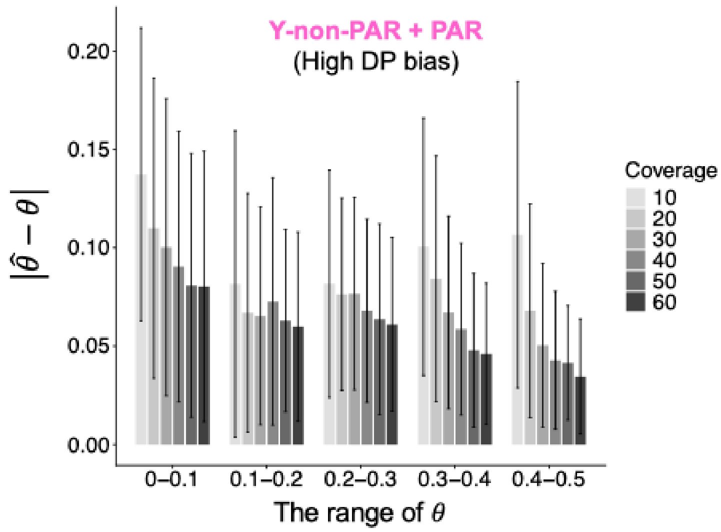


**Supplementary Figure S4.** Performance across different sequencing coverages and LOY ranges is shown for BaySeq-Y (Y-non-PAR + PAR) + using WES-like data ($n$ = 20 variants).

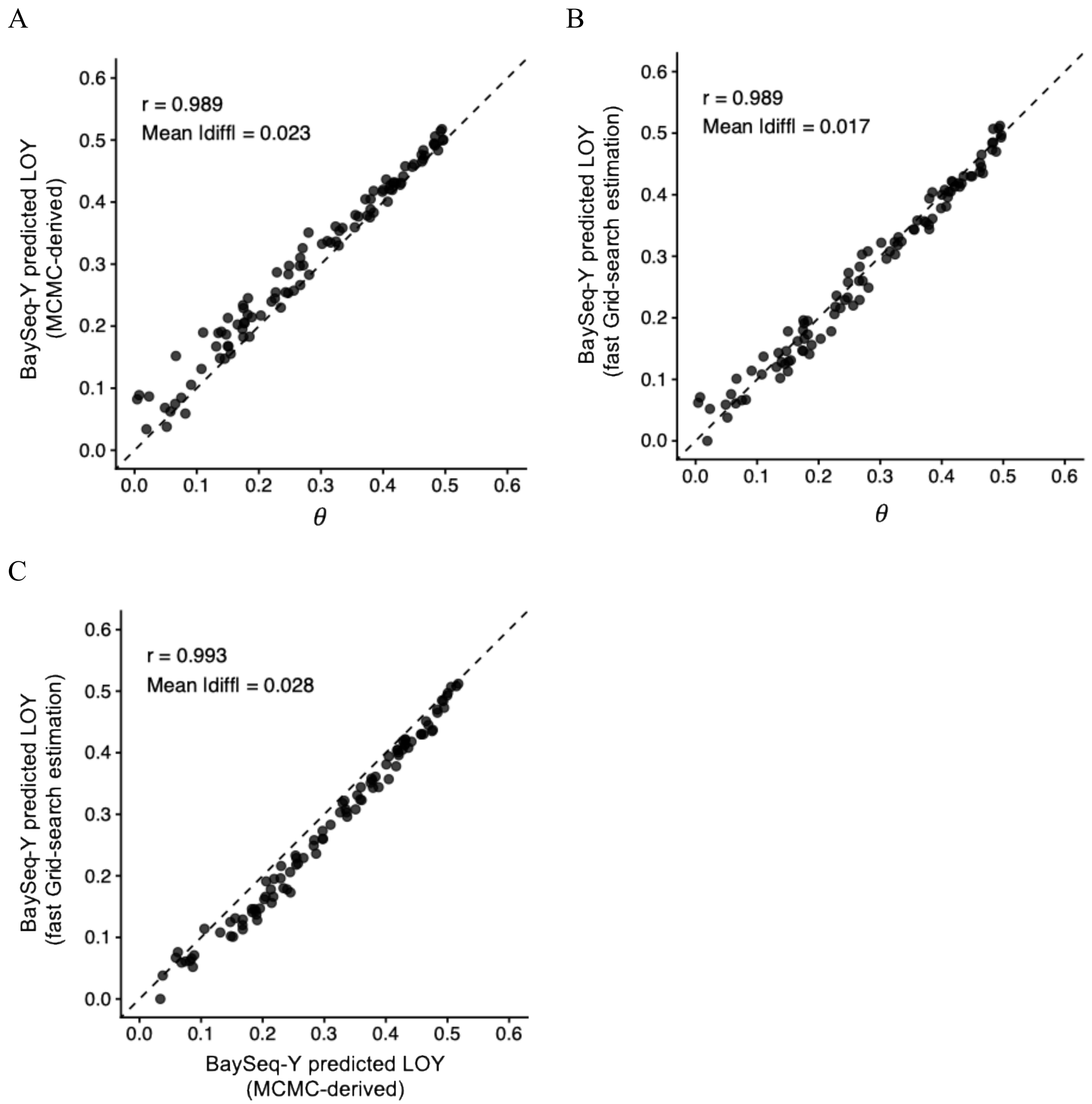


**Supplementary Figure S5.** MCMC-derived estimation vs. Grid-search estimation for BaysSeq-Y with phased data. Simulation-based performance of BaySeq-Y using phased PAR1 data with moderate phasing quality (see definition in **Methods**) was evaluated using different approximation algorithms for the Bayesian model.

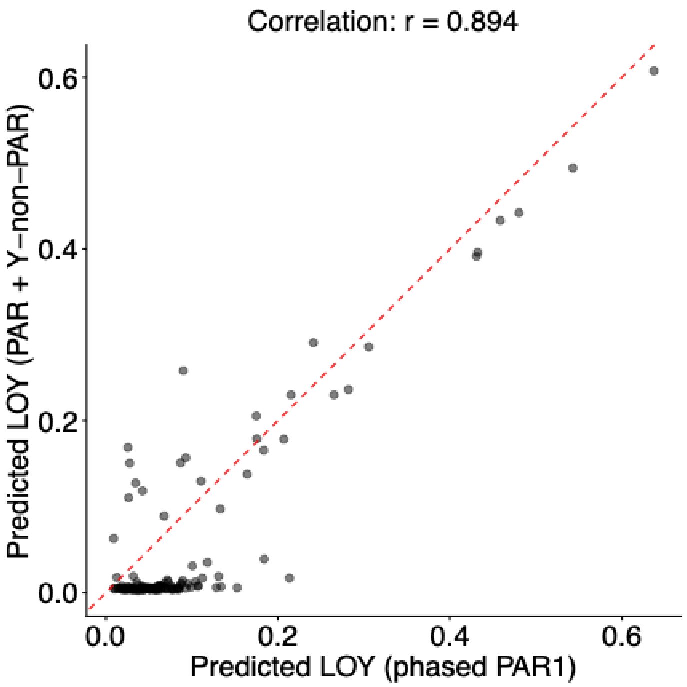


**Supplementary Figure S6.** Comparison of predicted LOY in ROSMAP samples using phased PAR1 and combined PAR/Y-non-PAR LOY-associated signals.

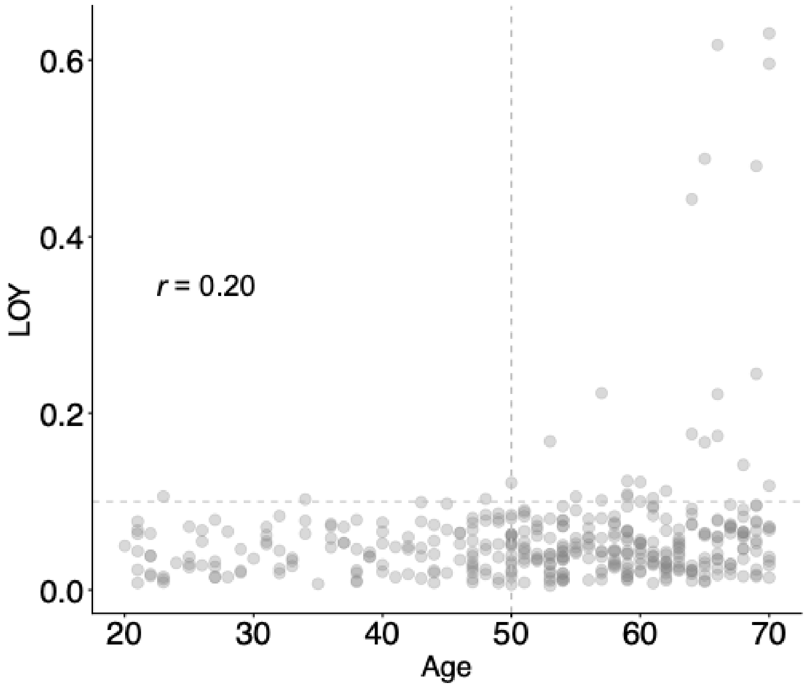


**Supplementary Figure S7.** LOY prediction in 377 GTEx males using BaySeq-Y with phased PAR1 data.

### Supplementary Tables

**Table S1.** Association between predicted LOY and whole-blood gene expression for PAR and Y-non-PAR genes in 256 GTEx males aged 50 years or older.

| Gene | P_value | FDR | Region in Y |
|---|---|---|---|
| ***UTY*** | 9.92E-29 | 2.08E-27 | Y non-PAR |
| ***DDX3Y*** | 1.16E-20 | 1.22E-19 | Y non-PAR |
| ***RPS4Y1*** | 5.67E-19 | 3.97E-18 | Y non-PAR |
| ***ZFY*** | 3.05E-13 | 1.60E-12 | Y non-PAR |
| ***KDM5D*** | 1.12E-09 | 4.71E-09 | Y non-PAR |
| ***EIF1AY*** | 1.26E-08 | 4.39E-08 | Y non-PAR |
| ***SLC25A6*** | 8.46E-08 | 2.54E-07 | PAR |
| ***DHRSX*** | 4.80E-06 | 1.26E-05 | PAR |
| ***CSF2RA*** | 5.96E-06 | 1.39E-05 | PAR |
| ***GTPBP6*** | 1.03E-04 | 2.17E-04 | PAR |
| ***TMSB4Y*** | 2.12E-04 | 3.78E-04 | Y non-PAR |
| ***P2RY8*** | 2.16E-04 | 3.78E-04 | PAR |
| ***ASMTL*** | 2.05E-03 | 3.32E-03 | PAR |
| ***USP9Y*** | 2.76E-03 | 4.14E-03 | Y non-PAR |
| ***PLCXD1*** | 1.39E-02 | 1.94E-02 | PAR |
| ***IL3RA*** | 2.40E-02 | 3.01E-02 | PAR |
| ***CD99*** | 2.43E-02 | 3.01E-02 | PAR |
| *AKAP17A* | 5.61E-02 | 6.54E-02 | PAR |
| *PPP2R3B* | 1.22E-01 | 1.35E-01 | PAR |
| *VAMP7* | 3.37E-01 | 3.54E-01 | PAR |
| *SPRY3* | 4.83E-01 | 4.83E-01 | PAR |
